# A membrane sensing mechanism couples local lipid metabolism to protein degradation at the inner nuclear membrane

**DOI:** 10.1101/2022.07.06.498903

**Authors:** Shoken Lee, Holly Merta, Jake W. Carrasquillo Rodríguez, Shirin Bahmanyar

## Abstract

Lipid composition is a determinant of organelle identity; however, whether the inner nuclear membrane (INM) domain of the endoplasmic reticulum (ER) harbors a unique lipid chemistry that contributes to its identity is not known. Here, we demonstrate that a unique INM lipid environment enriched in diacylglycerol protects the nucleo-cytoskeletal linker Sun2 from local degradation by the ubiquitin-proteasome system. A membrane binding amphipathic helix in the nucleoplasmic domain of Sun2 senses INM lipids and is essential to its protein stability. We show that the protein phosphatase CTDNEP1 localizes to the INM to maintain a distinct INM lipid environment necessary for Sun2 accumulation through regulation of the phosphatidic acid phosphatase lipin 1. Thus, the INM lipid environment sculpts the INM proteome via direct lipid-protein interactions that regulate protein stability, which has broad implications for mechanisms of diseases associated with the nuclear envelope.

## Introduction

The nuclear envelope (NE) is a highly specialized domain of the endoplasmic reticulum (ER) (Baumann and Walz, 2001; Hetzer, 2010). The NE serves as a selective permeability barrier to the genome and the ER is the site of synthesis for integral membrane proteins and for major membrane and storage lipids (Friedman and Voeltz, 2011). The bilayer lipids of the ER and NE encase a shared lumen and form distinct membrane structures: the ER is made up of small ribosome-bound sheets and membrane tubules while the NE is a large double membrane sheet (Friedman and Voeltz, 2011). The inner nuclear membrane (INM) faces chromatin and is adjoined with the outer nuclear membrane (ONM), which is directly continuous with the perinuclear ER. Nuclear pore complexes reside at fusion points between the INM and ONM to direct traffic of macromolecules across the NE (Ungricht and Kutay, 2017).

The segregation of the INM from the cytoplasm provides a confined environment for the NE to carry out its unique functions, which include nucleo-cytoskeletal coupling, genome organization and lipid metabolism (Bahmanyar and Schlieker, 2020; Starr and Fridolfsson, 2010; Ungricht and Kutay, 2017). As such, the INM harbors a unique proteome. In metazoans, the intermediate filament lamin proteins form a meshwork at the nuclear face of the INM to provide mechanical stability to the NE (Dechat et al., 2010; Dechat et al., 2008). A subset of integral membrane proteins diffuse from the site of synthesis on ribosome-bound ER sheets to the INM where they selectively concentrate through interactions with chromatin and the nuclear lamina (Boni et al., 2015; Katta et al., 2014; Ungricht et al., 2015). Mutations in nuclear lamins and INM proteins give rise to a diverse array of human disorders including muscular dystrophies, lipodystrophy, and progeria, further highlighting the importance of INM functions to cellular and human physiology (Shin and Worman, 2022). Both active transport and passive diffusion of membrane proteins via the NPC are important to establish the INM proteome (Katta et al., 2014); however, how cells adapt and remodel the INM proteome in response to changes to cellular conditions is not well understood.

Some evidence also suggests that dedicated protein degradation mechanisms selectively monitor and refine the INM proteome. In budding yeast, an E3 ligase complex selectively targets unwanted ER proteins for degradation at the INM (Foresti et al., 2014; Khmelinskii et al., 2014; Natarajan et al., 2020). In mammalian cells, exogenously expressed INM protein disease mutants are locally targeted for degradation by the proteasome (Tsai et al., 2016) and can be cleared from the INM via vesicular transport to the lysosome (Buchwalter et al., 2019). Local protein turnover of resident INM proteins could serve to dynamically control the steady state profile of INM proteins. However, whether wild type resident INM proteins are also susceptible to regulated proteolysis at the INM has not been shown.

Direct protein binding to a unique INM lipid environment (e.g., lipids with specific head groups or lipid packing density) could provide a way to target, enrich and remodel the INM proteome. In the vesicular trafficking pathway, the unique bilayer lipid composition of membrane-bound compartments confers organelle identity by recruiting proteins with specific lipid binding domains. Lipids can be quickly metabolized by lipid modifying enzymes allowing for rapid changes in organelle identity (Behnia and Munro, 2005). It is likely that similar mechanisms exist at the INM; however, little is known about how a unique INM lipid composition is established and maintained despite the possibility of free lipid diffusion with continuous ER membranes. Key findings in budding yeast and mammalian cells revealed that INM lipids are dynamically altered (Barbosa et al., 2019; Haider et al., 2018; Romanauska and Köhler, 2018, 2021; Sołtysik et al., 2021; Tsuji et al., 2019); however, it is unclear if INM-resident proteins respond to lipid metabolism at the INM. In budding yeast, protein biosensors appended to nuclear localization signals revealed enrichment of diacylglycerol (DAG) at the INM at steady state (Romanauska and Köhler, 2018). Protein biosensors that detect the steady state content of INM lipids in mammalian cells have not previously been developed.

Here, we establish a protein biosensor that detects DAG at the INM in mammalian cells. DAG enrichment at the INM depends on the phosphatidic acid phosphatase lipin 1 (Pah1p in budding yeast/Ned1 in fission yeast/LPIN-1 in C. elegans) (Kim et al., 2007; O’Hara et al., 2006; Tange et al., 2002) and its activating protein phosphatase, CTDNEP1 (Nem1 in budding and fission yeasts/CNEP-1 in *C. elegans*) (Bahmanyar et al., 2014; Han et al., 2012; O’Hara et al., 2006; Santos-Rosa et al., 2005; Siniossoglou et al., 1998), which we show localizes to the INM together with its binding partner NEP1R1. We find that the protein levels of the nucleo-cytoskeletal linker Sun2 is sensitive to alterations of INM DAG content. An amphipathic helix (AH) of Sun2 confers this sensitivity to INM lipids and is also essential for stability of Sun2 protein. Furthermore, we demonstrate that a cluster of serine residues directly upstream of the membrane sensing AH of Sun2 determines its protein stability. Live imaging of NE-associated Sun2 in cells acutely depleted of CTDNEP1 revealed that Sun2 loss occurs locally at the INM in response to metabolic changes in INM lipids. Thus, preferential membrane binding of Sun2 to INM lipids protects against its targeting for protein degradation. Our data suggest a model in which active mechanisms maintain the unique composition of INM lipids relative to the ER to sculpt the INM proteome and thus maintain its unique functions in genome regulation and protection.

## Results

### Local control of diacylglycerol levels revealed by an INM-DAG biosensor

Lipid metabolism occurs locally at the INM but mechanisms that directly control the bilayer lipid content of the INM at steady state are not fully understood. We chose to focus on the NE-enriched protein phosphatase CTDNEP1 because of its known role for regulating the phosphatidic acid phosphatase lipin 1 (Figure 1A) (Han et al., 2012; Kim et al., 2007; Merta et al., 2021), which is mostly soluble at steady state and localizes to both the nucleus and cytoplasm (Merta et al., 2021; Peterson et al., 2011). We used CRISPR-Cas9 genome editing to endogenously tag human CTDNEP1 with EGFP at the endogenous locus (CTDNEP1(EN)-EGFP; Figure 1B). CTDNEP1(EN)-EGFP localized to the NE in U2OS cells, as well as to other cytoplasmic membrane structures that likely represent an ER-associated pool of the endogenous protein (Figure 1B). The split-sfCherry2 system (Feng et al., 2017) in which a piece of sfCherry_1-10_ is fused to either CTDNEP1 or NEP1R1 was co-transfected with sfCherry_11_ fused to Histone-2B revealed the population of CTDNEP1 and NEP1R1 that can reach the INM (Figure 1C). Localization of both CTDNEP1 and NEP1R1 to the INM suggested that the INM lipid environment, and particularly PA/DAG metabolism, in animal cells is locally controlled through lipin 1 regulation.

**Figure 1:**
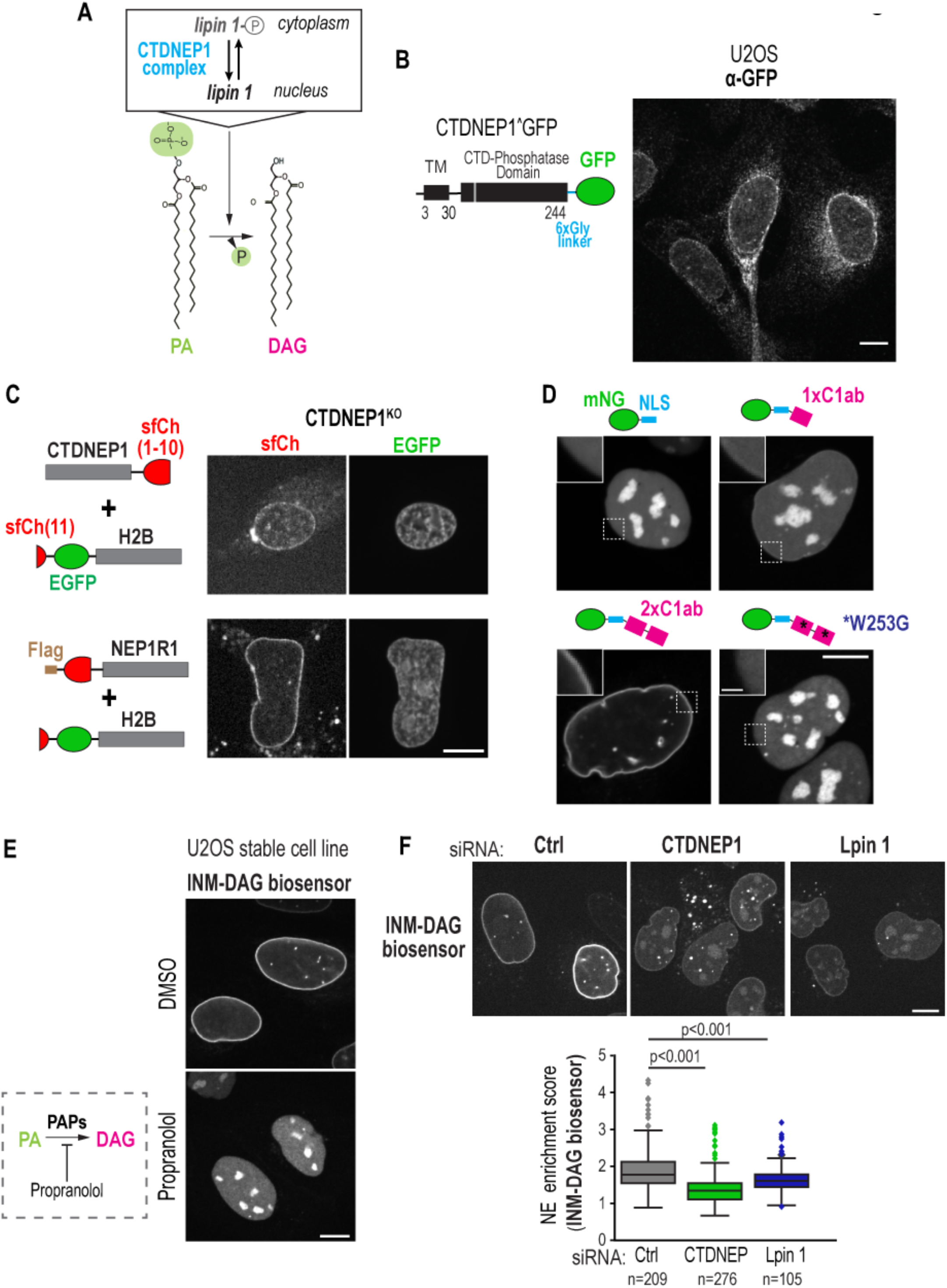
CTDNEP1 and lipin 1 control enrichment of diacylglycerol at the inner nuclear membrane. (A) Schematic representation of CTDNEP1 and lipin 1 in relation to production of diacylglycerol. (B) Confocal images of CTDNEP1(EN)-GFP in U2OS cells immunostained with anti-EGFP. (C) Spinning disk confocal images of live U2OS cells expressing indicated constructs. (D) Spinning disk confocal images of live U2OS cells expressing the indicated DAG biosensors. (E) Spinning disk confocal images of live U2OS cells stably expressing the INM-DAG biosensor and treated with propranolol. (F) Spinning disk confocal images of live U2OS cells stably expressing the INM-DAG biosensor and treated with indicated siRNAs. Box plot, NE enrichment scores pooled from 3-4 independent experiments. p values, Tukey HSD test. Scale bars, 2 µm in zoom in (D), 10 µm in others.

We established genetically encoded protein biosensors to detect the DAG pool at the INM by utilizing the C1a-C1b (hereafter C1ab) domains of protein kinase C theta, which had been used for detecting DAG at other organelles (Carrasco and Merida, 2004; Spitaler et al., 2006). A single C1ab domain appended to an NLS showed faint localization at the nuclear rim, whereas a tandem repeat of the C1ab domains (2xC1ab) fused to an NLS (hereafter referred to as INM-DAG biosensor) showed clear nuclear rim localization with intranuclear localization that resembles nucleoplasmic reticulum (Figure 1D). The INM-DAG biosensor with its key residue for DAG binding mutated (W253G) (Das and Rahman, 2014; Rahman et al., 2013) localized to the nucleoplasm and nuclear bodies in a non-specific manner that was indistinguishable from localization of mNG-NLS indicating the requirement for DAG binding for the localization of the biosensor at the INM (Figure 1D). Localization of the INM-DAG biosensor to the nuclear rim was abolished after 5 minutes in cells treated with a small molecule inhibitor of Mg^2+^-dependent and -independent PA phosphatases (propranolol; Figure 1E), which has been shown to deplete DAG from cytosolic organelle membranes (Baron and Malhotra, 2002; Carrasco and Merida, 2004). RNAi-mediated depletion of CTDNEP1 and lipin 1 after 72 hours showed a reduction of the INM-DAG biosensor from the nuclear rim, albeit to a lesser extent than short-term propranolol treatment, and concomitant localization to puncta outside the nucleus that likely represent a pool of DAG at cytoplasmic organelles (Figure 1F). To quantitatively confirm the reduced levels of the INM-DAG biosensor at the nuclear rim in an unbiased manner, we used semi-automated image analysis to determine the ratio of the fluorescent signal of the INM-DAG biosensor at the nuclear rim and nucleoplasm (‘NE enrichment score’; Figure S1). This analysis further confirmed that NE enrichment of the INM-DAG biosensor, and thus DAG levels at the INM, requires CTDNEP1 and lipin 1 (Figure 1F).

### Reduction of INM DAG levels corresponds to loss of the nucleo-cytoskeletal linker Sun2

We reasoned that, similar to cytoplasmic organelles, the unique lipid environment of the NE plays a role in establishing or maintaining the identity of the NE through regulation of its protein composition. We compared the endogenous nuclear rim localization of six INM proteins in control U2OS and CRISPR-Cas9 edited CTDNEP1 KO cells (Merta et al., 2021) and found a dramatic reduction in fluorescence signal specific to Sun2 (Figure S2A; Figure 2A). The reduction of Sun2 in CTDNEP1 KO cells was restored upon expression of wild type CTDNEP1 (CTDNEP1-mAID-HA), but not the phosphatase dead mutant (phosphatase-dead, PD) (Figures 2A and 2B) (Merta et al., 2021). Sun proteins are a family of nucleo-cytoskeletal linkers defined by a luminal SUN domain at their C-terminus that facilitates binding to the cytoskeleton through direct association with KASH proteins in the ONM (Chang et al., 2015). Sun1 and Sun2 are ubiquitously expressed and recent studies indicate that Sun2 has cellular and tissue-level functions distinct from Sun1 (Luxton et al., 2010; Stewart et al., 2019; Stewart et al., 2015; Zhu et al., 2017). The selective reduction of Sun2 in CTDNEP1 KO cells suggests a potential regulatory mechanism specific to Sun2 function that involves INM lipid metabolism.

**Figure 2:**
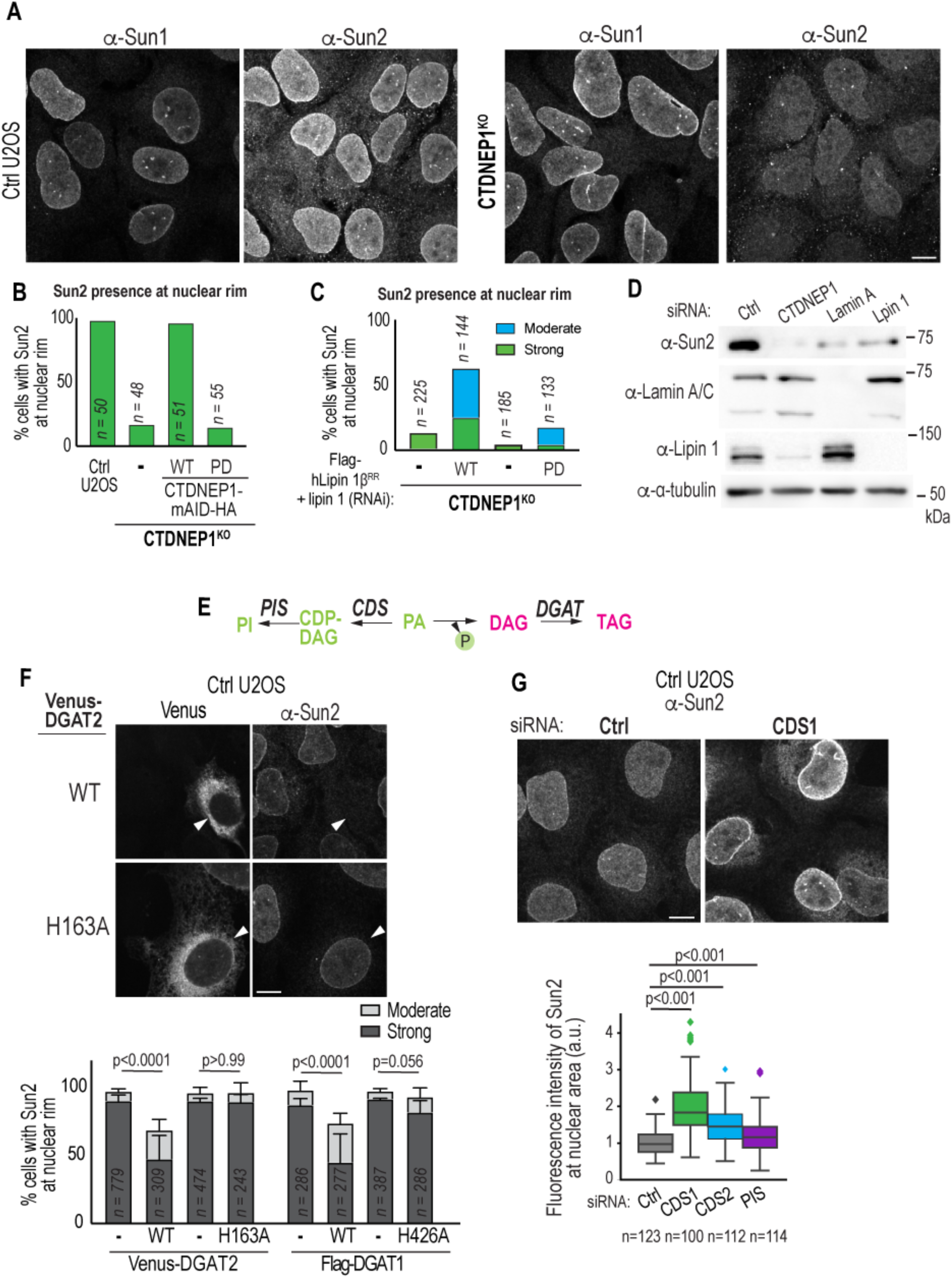
Alterations in lipid metabolism centered on DAG affect Sun2 localization and protein levels. (A) Confocal images of immunostained U2OS cells for indicated proteins. (B and C) Plot of quantification of Sun2 localization at nuclear rim. (D) Immunoblot of control U2OS cells treated with indicated siRNAs. (E) A simplified schematic of lipid synthesis pathway in metazoans. (F) Confocal images of immunostained U2OS cells expressing indicated constructs. Plot shows quantification of Sun2 localization at nuclear rim. N = 3 independent experiments. Mean + SD shown. p values, Fisher’s exact test (Clear vs others). (G) Confocal images of immunostained U2OS cells treated with indicated siRNAs. Box plot, Sun2 fluorescence intensity in nuclear area. p value, Tukey HSD test. Scale bars, 10 µm

Importantly, overexpression of wild type human lipin 1, but not the catalytically inactive mutant, restored the localization of Sun2 at the nuclear rim in CTDNEP1 KO cells RNAi-depleted of endogenous lipin 1, as shown by blind categorization (Figure 2C) and unbiased automated image analysis (Figure S2B). We found a global reduction of Sun2 protein and not transcript levels in CTDNEP1 KO cells (Figure 2D; Figure S2C). Lower lipin 1 protein levels result from RNAi-depletion of CTDNEP1 (Figure 2D), as expected (Merta et al., 2021), but this does not affect lamin A/C protein levels or Sun1 localization at the NE, supporting the possibility that general NE defects are unlikely to be the cause of Sun2 loss (Figure 2D; also see Figures S2D and S2E). RNAi-depletion of lipin 1 also showed a reduction in global Sun2 protein levels (Figure 2D) and loss of Sun2 localization at the nuclear rim (Figure S2D) similar to loss of CTDNEP1. The effect of RNAi-depletion of lipin 1 was slightly less pronounced than that from depletion of CTDNEP1 (Figure 2D), possibly due to differences in knockdown levels or compensatory mechanisms that arise upon loss of lipin 1 (Grimsey et al., 2008). Together, these results indicate that Sun2 levels are controlled by the lipid modifying activity of lipin 1, which is known to be regulated and stabilized by NE-enriched CTDNEP1 (Han et al., 2012; Kim et al., 2007; Merta et al., 2021; Zhang and Reue, 2017).

We hypothesized that the effects on Sun2 resulting from loss of CTDNEP1 are because of local changes in INM lipids and not global upregulation of *de novo* lipid synthesis that we previously reported occurs in these cells (Merta et al., 2021). Indeed, the nuclear rim localization of Sun2 in CTDNEP1 KO cells was unaffected in cells treated with TOFA, a small molecule inhibitor of the rate limiting enzyme in *de novo* fatty-acid synthesis that restores ER structure in these cells (Figure S2F) (Merta et al., 2021). Furthermore, the fact that we did not observe reduction of emerin in CTDNEP1 KO cells (Figure S2A), which has been shown to be downregulated along with Sun2 upon disruption of ER homeostasis (Buchwalter et al., 2019), further indicates that a selective mechanism by CTDNEP1 control of lipin 1 regulates Sun2 protein levels. Sun2 is anchored to the NE by nuclear lamins (Chang et al., 2015), however we also did not observe a change in lamin A/C levels upon loss of CTDNEP1 or lipin 1 (Figure 2D). Lamin A/C knockdown caused nonselective mislocalization of both Sun1 and Sun2 (Figures S2D and S2E) further indicating that CTDNEP1/lipin 1 likely control Sun2 levels independently of lamin A/C. These data supported our hypothesis that lipid metabolism-related effects resulting from loss of CTDNEP1/lipin 1 are likely directly responsible for the changes in Sun2 protein levels.

We next tested if manipulating other lipid metabolic enzymes in the *de novo* lipid synthesis pathway that are predicted to affect DAG consumption or production would alter Sun2 localization at the nuclear rim in wild type cells (Figure 2E). DGAT1 and DGAT2 can localize to the INM (Sołtysik et al., 2021) and consume DAG to generate triglycerides (TAG) (Figure 2E). A significant proportion of cells overexpressing DGAT1 and DGAT2 showed loss of Sun2, but not Sun1, at the nuclear rim (Figure 2F; Figures S3A and S3B). Importantly, overexpression of catalytically inactive DGAT1 and DGAT2 mutants (McFie et al., 2010; Stone et al., 2006) did not affect Sun2 levels at the nuclear rim, supporting the idea that Sun2 is sensitive to DAG consumption by these enzymes. Furthermore, there was an increase in Sun2 levels at the nuclear rim resulting from RNAi-depletion of CDS1, CDS2 or PIS, which are predicted to increase flux towards PA conversion to DAG by reducing PA flux towards PI synthesis (Bahmanyar et al., 2014) (Figure 2G; Figure S3C). Taken together, we conclude that consumption or production of DAG at the INM regulates Sun2 accumulation at the nuclear rim.

### Characterization of a membrane binding amphipathic helix in Sun2

Next, we aimed to identify a region in Sun2 that directly senses the INM lipid environment. Exogenously overexpressed Sun2-mNeonGreen (mNG), but not Sun1-mNG, results in a reduced NE enrichment and increased ER localization in CTDNEP1 KO cells (Figure S4A) further indicating that a unique feature in Sun2, that is not present in Sun1, is responsive to metabolic changes in INM lipids resulting from loss of CTDNEP1. Sun1 and Sun2 both contain an N-terminal nucleoplasmic (NP) domain, followed by hydrophobic alpha helices (Liu et al., 2007; Majumder et al., 2018; Turgay et al., 2010), a transmembrane helix (or helices in Sun1 (Majumder et al., 2018)), and a large luminal domain that oligomerizes and spans the luminal space (Sosa et al., 2012). Swapping the transmembrane and luminal domains of Sun2 with those of Sun1 (Sun2(1-209)-Sun1(220-716)) phenocopied the reduced enrichment of full length Sun2-mNG at the NE of CTDNEP1 KO cells (Figure S4B). In contrast, a chimera in which the NP domain of Sun2 is swapped with that of Sun1 (Sun1(1-219)-Sun2(213-717)) caused enrichment at the NE to the same extent as full length Sun1-mNG (Figure S4B). These data suggested that the NP region of Sun2, but not of Sun1, harbors an amino acid sequence that renders Sun2 sensitive to the lipid environment. Several previous studies have addressed the significance of the NP region of Sun2 in its targeting to the nuclear rim (Crisp et al., 2006; Hasan et al., 2006; Turgay et al., 2010; Wang et al., 2006); however, these studies did not test the specific contribution of the hydrophobic helices in this region.

The Sun1 and Sun2 lamin-binding region resides at the very N-terminus of the NP domain (Haque et al., 2010) (Figure S4C). Swapping the lamin-binding region of Sun2 with that of Sun1 was sufficient for the Sun1-Sun2 chimera to enrich at the NE (Sun1(1-136)-Sun2(126-717)) in CTDNEP1 KO cells (Figure S4C). Thus, the lamin-binding region of Sun1 retains Sun1-Sun2 chimeras at the NE and overrides any sensitivity in the NP domain of Sun2 to the INM lipid environment. These data highlight that different mechanisms enrich Sun1 and Sun2 at the NE.

Our *in silico* analysis predicted the presence of an amphipathic helix (AH) (residues 155-180) in the Sun2 NP domain downstream the N-terminal lamin binding region (Figure 3A). A series of chimeras between Sun1 and Sun2 were used to further test if the predicted AH in Sun2 confers its sensitivity to loss of CTDNEP1. A Sun1-Sun2 chimera with the Sun2 AH (Sun2(1-180)-Sun1(191-716)) reduced enrichment at the NE in CTDNEP1 KO cells, whereas a chimera that does not contain the Sun2 AH (Sun2(1-150)-Sun1(156-716)) retained enrichment at the NE (Figure S4D). These data suggested that the predicted AH in Sun2 is a key determinant that renders Sun2 sensitive to the NE lipid environment.

**Figure 3:**
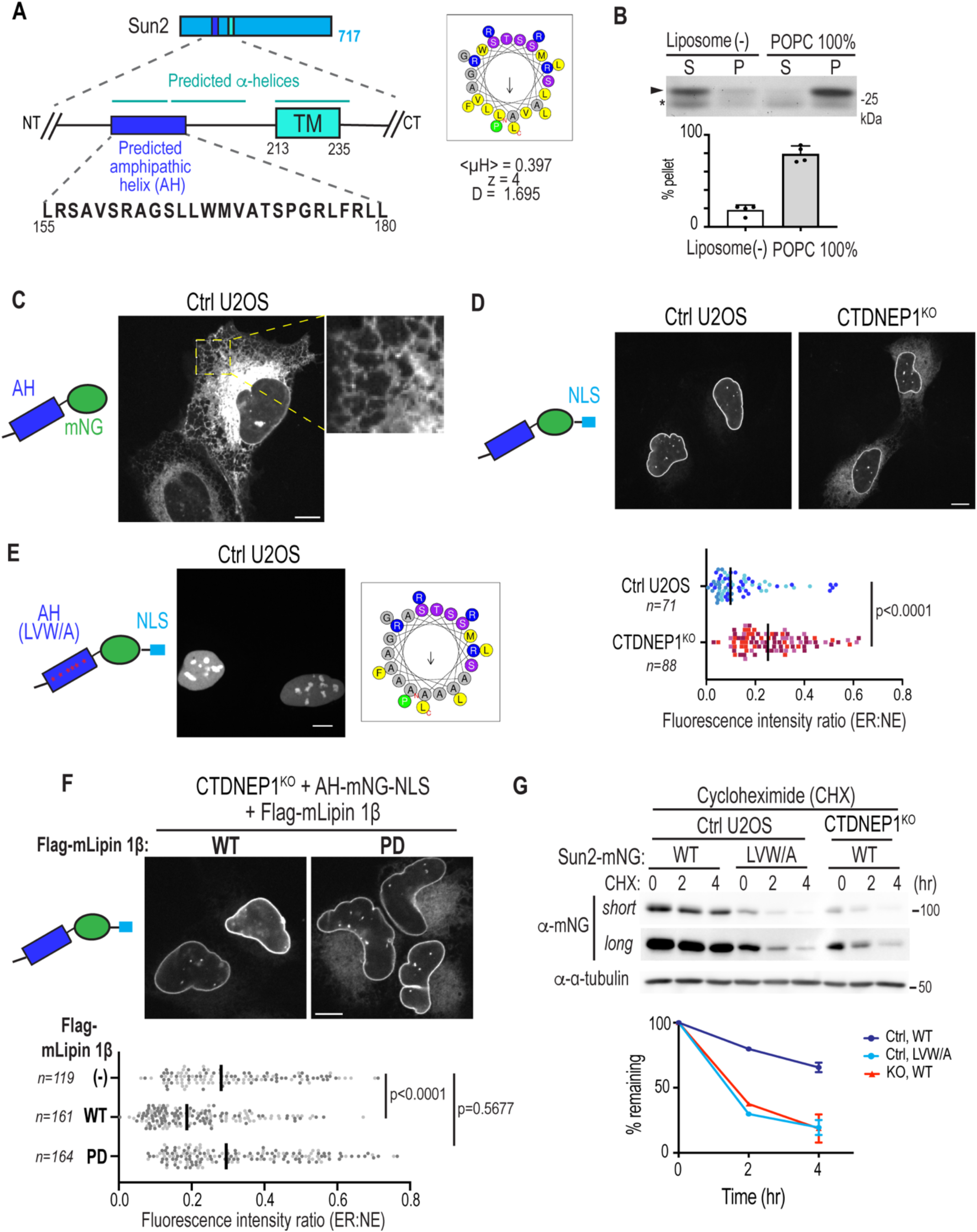
Sun2 contains a membrane binding amphipathic helix that controls its protein levels and is sensitive to the INM lipid environment. (A) Schematic of N-terminal region of Sun2 highlighting predicted amphipathic helix (residues 155-180). A helical wheel projection generated by HeliQuest with hydrophobic moment <µH>, net charge z, and the discriminant factor D. (B) Liposome-cosedimentation with recombinant GST-tagged AH peptide (residues 151-180) and 1-palmitoyl-2-oleoyl-sn-glycero-3-phosphocholine (POPC) liposome. Plot, percent of protein found in pellet as a ratio of soluble fraction. Four experimental replicates. Arrowhead, GST-AH; Asterisk, purification biproduct. (C-E) Spinning disk confocal images of U2OS cells expressing indicated constructs. Plot in (D) shows ER:NE fluorescence ratio pooled from 3 independent experiments. p value, Mann-Whitney U test. In (E), a helical wheel projection projection generated by HeliQuest of AH(LVW/A) mutant. (F) Spinning disk confocal images of U2OS cells expressing AH-mNG-NLS and Flag-mLipin 1β. Plot, ER:NE fluorescence ratio pooled from 2 independent experiments. p values, ordinary one-way ANOVA followed by Dunnett’s multiple comparisons test. (G) Cycloheximide chase assay of cells expressing indicated constructs. N = 2 (2 hr) or 3 (4 hr) independent experiments. Mean (and SD where N = 3) shown. In (D) and (F), bars indicate median, dots are color coded according to experimental replicates. Scale bars, 10 µm

We hypothesized that the predicted AH of Sun2 may directly bind and sense the INM lipid environment. Not much is known about the specific bilayer lipid make-up of the INM and so we tested if this region binds directly to lipid bilayers with a preference for a specific membrane property that may resemble the INM enriched in DAG. We found that a peptide with the Sun2 AH (residues 151-180) purified from *E. coli* as a GST fusion binds to palmitoyl-oleoyl-phosphatidylcholine (POPC) liposomes *in vitro* (Figure 3B), whereas a PS-specific PH domain (Uchida et al., 2011) does not (Figure S5A), indicating that this portion of Sun2 contains a direct membrane binding helix. DAG is known to induce packing defects in lipid bilayers (Vamparys et al., 2013; Vanni et al., 2014); however, our attempts to generate liposomes with DAG were unsuccessful, possibly due to membrane-fusogenic effect of DAG observed *in vitro* (Gómez-Fernández and Corbalán-García, 2007). Instead we turned to generating liposomes with saturated acyl chains or cholesterol, a method that is more commonly used (Kassas et al., 2017; Vanni et al., 2014), to test if increased lipid packing affected Sun2 AH binding. Indeed, increased lipid packing of liposomes resulted in decreased binding of the AH of Sun2 (Figures S5B-S5D). Together, these data demonstrate that the nucleoplasmic AH of Sun2 binds directly to membrane bilayers with a preference for reduced lipid packing. Thus, the AH of Sun2 may be more sensitive to bulk membrane properties that exist at the INM (such as membrane packing defects or fluidity) rather than the concentrations of specific lipid species (such as DAG).

Our *in vitro* data predicted that the AH of Sun2 can localize to ER/NE membranes through peripheral membrane binding when expressed in cells. The AH of Sun2 targeted to the ER when expressed from a plasmid as an mNG fusion in control U2OS cells (Figure 3C). Selective permeabilization of the plasma membrane versus internal membranes by digitonin (Adam et al., 1990) followed by antibody staining against the N-terminal HA- and C-terminal mNG tags appended to residues 151-180 of Sun2 demonstrated that the AH does not cross the membrane bilayer (Figure S6A). Importantly, Sun2-AH appended with an NLS (hereafter AH-mNG-NLS) was strongly enriched at the NE in control U2OS cells (Figure 3D). Furthermore, mutating six of the eleven large hydrophobic residues in the AH of AH-mNG-NLS (hereafter LVW/A mutant) completely abolished its localization at the INM (Figure 3E). Together, these data indicate that the AH of Sun2 is a membrane-associated amphipathic helix that enriches at the INM when targeted to the nucleus.

The lipid-binding AH of Sun2 is sufficient to confer sensitivity to changes in INM lipid metabolism caused by CTDNEP1 (Figure 3D). We observed loss of NE enrichment in AH-mNG-NLS in CTDNEP1 KO cells concomitant with its dispersal to the ER (Figure 3D), which was rescued by expression of wild type CTDNEP1, but not the phosphatase-dead version (Figure S6B). The dispersed localization of AH-mNG-NLS was partially restored by overexpression of lipin 1, but not the phosphatase inactive mutant, in CTDNEP1 KO cells (Figure 3F) suggesting that the NE enrichment of the amphipathic helix of Sun2 depends on INM lipid identity centered on PA-DAG metabolism. The dispersal of the AH to the ER, rather than to the nucleoplasm, in CTDNEP1 KO cells suggested that disruption of the unique INM lipid environment leads to the indiscriminate association of AH-mNG-NLS to ER membranes. We cannot exclude the possibility that other defects, such as loss of NE integrity or defective nuclear transport, affects the steady state enrichment of AH-mNG-NLS at the INM in CTDNEP1 KO cells, although an mNG-NLS reporter was normally retained in the nucleus in these cells (Figure S6C).

Thus, we conclude that a direct membrane binding by the AH in the nucleoplasmic region of Sun2 confers specificity for localization to the unique INM lipid environment, which is at least in part maintained by CTDNEP1/lipin 1.

### Sun2 protein stability depends on its membrane-binding amphipathic helix

Prior work has shown that Sun2 is targeted for degradation by the proteasome (Ji et al., 2022; Kim et al., 2015; Loveless et al., 2015). A degradation mechanism triggered by reduced INM association of the AH of Sun2 would explain the lower steady state levels of full length Sun2 protein in CTDNEP1 KO cells (see Figure 2D). Cycloheximide chase experiments showed that Sun2-mNG is short-lived in CTDNEP1 KO cells when compared to control U2OS cells (Figure 3F). Importantly, the mutant Sun2-mNG protein that no longer binds to the INM (AH-LVW/A) lost its enrichment at the NE in control U2OS cells (Figure S6D) and is short-lived with degradation kinetics similar to that of wild type Sun2-mNG in CTDNEP1 KO cells (Figure 3F). These data indicate that the lipid-binding AH is part of a degradation mechanism that controls the half-life of Sun2 and that hampering INM lipid association by Sun2, either by mutating the hydrophobic residues of its AH helix or by altering the INM lipid environment through loss of CTDNEP1, causes protein destabilization.

### Proteasomal degradation of Sun2 is coupled to the membrane binding amphipathic helix of Sun2

We sought to more directly determine if INM dissociation of the Sun2 AH triggers its degradation by the proteasome. Degradation of Sun2 is mediated by the substrate recognition subunit of the Skp-Cullin-F-box (SCF), beta-transducin repeats-containing proteins b-TrCP1/Fbxw1 and b-trcp2/Fbx11, which are Cullin-Ring ligases referred to SCF^b-trcp^ (Kim et al., 2015; Loveless et al., 2015). We found that the steady state levels of endogenous Sun2 in CTDNEP1 KO cells were partially restored either in cells treated with a small molecule inhibitor against the Nedd8-activating enzyme (MLN4924) that inactivates Cullin-Ring ligases (Figure 4A) or with RNAi-depletion of SCF^b-trcp^ subunits (Figure S7A). A cycloheximide chase experiment showed that Sun2-mNG is longer-lived in CTDNEP1 KO cells treated with MLN4924 in which SCF^b-trcp^ function was inhibited (Figure S7B). These data confirm that Sun2 is likely targeted for proteasomal degradation by SCF^b-trcp^.

**Figure 4:**
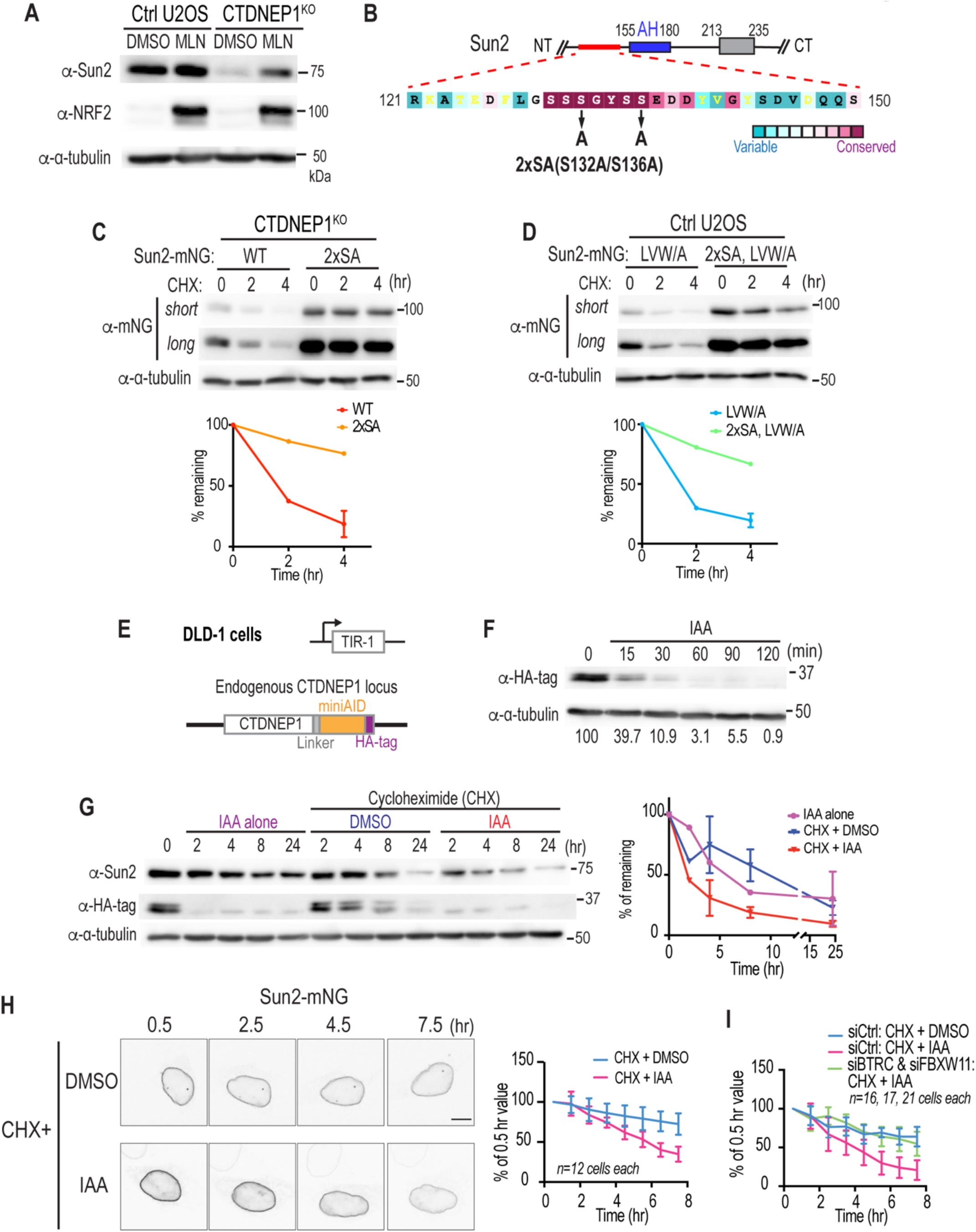
Requirements for Sun2 protein stability and loss of a NE-associated pool of Sun2 via the proteasome. (A) Immunoblot of indicated cells treated with MLN4924. (B) Human Sun2 residues 121-150 color-coded according to ConSurf conservation scores of vertebrate Sun2 protein sequences. (C and D) Cycloheximide chase assay of indicated cells expressing indicated constructs. Plots of wild-type and LVW/A are replicas of Figure 3F. N = 2 or 3 (wild type and LVW/A, 4 hr) independent experiments. Mean (and SD where N = 3) shown. (E) Schematic of CTDNEP1(EN)-mAID-HA. (F and G) Immunoblot of DLD-1 OsTIR-1 CTDNEP1(EN)-mAID-HA cells treated as indicated. In (E), values at the bottom show quantification of blot of HA-tag, normalized to α-tubulin blot and shown as % of 0 min. In (G), N = 2 (IAA alone and all 2 hr) or 3 independent experiments. Mean (and SD where N = 3) shown. (H and I) Time-lapse confocal images of transiently expressing Sun2-mNG in live DLD-1 CTDNEP1(EN)-mAID-HA cells stably expressing OsTIR-1, treated as indicated. mNG intensity at nuclear rim is shown as % of the value at 0.5 hr. Data were pooled from 2 independent experiments. Scale bar, 10 µm.

We performed amino acid alignments of vertebrate Sun2 to search for a consensus degron sequence that may be recognized directly by SCF^b-trcp^. We identified a highly conserved serine cluster (aa131-136; SSGYSS) directly upstream of the Sun2-AH with some resemblance to the known SCF^b-trcp^ degron sequence (DSGXXS) (Frescas and Pagano, 2008) (Figure 4B). Expression of a mutant version of Sun2 in which two key serine residues were mutated to alanine (2xSA) led to its stabilization (Figure 4C) and restored its accumulation at the NE in CTDNEP1 KO cells (Figure S7C). We predicted that mutating the serine residues in Sun2 would recover its protein instability even in the absence of membrane binding through its AH. In line with this, the Sun2-AH(LVW/A) mutant, which is short-lived in control U2OS cells, was longer-lived with the addition of the 2xSA mutation in the upstream degron sequence (Figure 4D). Together, these data suggest a working model in which dissociation of the AH from the INM triggers proteasomal degradation of Sun2 through a neighboring sequence (‘Ser- cluster’) that may serve as a degron recognized by a nuclear pool of SCF^b-trcp^. The fact that Sun2 can be stabilized at the INM when degradation is inhibited (such as in the case of mutating the Sun2 degron) suggests Sun2 enrichment at the INM requires AH membrane-binding to prevent its degradation.

### The nuclear envelope associated pool of Sun2 is targeted for proteasomal degradation in response to changes in local lipid composition

To test if the INM pool of Sun2 is targeted for degradation, which would fit with our model that direct sensing of INM lipids by the AH of Sun2 is responsible for its stabilization, we turned to the auxin-inducible degron (AID) system to acutely deplete CTDNEP1 and determine the degradation kinetics of Sun2 protein (Natsume et al., 2016; Nishimura et al., 2009). We tagged endogenous CTDNEP1 with the mAID degron followed by an HA tag in DLD-1 cells stably expressing the OsTIR-1 F-box protein, which is required for auxin-dependent mAID recognition (Figure 4E). Treatment with auxin (indole-3-acetic acid, IAA) induced rapid degradation of CTDNEP1(EN)-mAID-HA within 1-2 hr (Figure 4F) and led to instability of endogenous Sun2 protein levels (Figure 4G). To determine if Sun2 loss occurs locally at the INM, we performed a cycloheximide chase experiment to monitor the levels of existing Sun2-mNG at the nuclear rim by live imaging following acute depletion of CTDNEP1. Sun2-mNG showed a slow loss of fluorescence signal at the nuclear rim in control cells under normal conditions, which may partially reflect its normal protein turnover rate. Acute depletion of CTDNEP1 accelerated the loss of Sun2-mNG fluorescence signal at the nuclear rim (Figure 4H). RNAi-depletion of SCF^b-trcp^ stabilized NE-associated Sun2-mNG in cells acutely depleted of CTDNEP1 (Figure 4I; Figures S7D and S7E). We also monitored the ER-associated pool of Sun2-mNG and found similar trends in the rate of loss of fluorescence (Figure S7F). The loss of Sun2-mNG fluorescence signal from the NE and its dependence on SCF^b-trcp^ suggests that the INM-associated pool of Sun2 is a target for proteasomal degradation. The fact that the ER-associated pool of Sun2-mNG is also unstable upon loss of CTDNEP1 suggests that Sun2 loss occurs throughout the ER and NE when lipid composition is not tightly regulated, possibly because of weakened membrane association via its AH.

## Discussion

Our data suggest a model in which a membrane-sensing amphipathic helix restricts the accumulation of the nucleo-cytoskeletal linker Sun2 to the INM (Figure 5). Several regions of Sun2 have been shown to play a role in its targeting to the INM (Haque et al., 2010; Turgay et al., 2010). We provide evidence that targeting and accumulation of Sun2 at the INM is mediated by direct membrane binding coupled to regulated proteolysis. We suggest that the amphipathic helix (AH) of Sun2 facilitates strong membrane association at the INM via direct lipid binding, which in turn mediates a conformational change that shields the upstream “serine cluster” to prevent targeting of Sun2 for degradation by the proteasome (Figures 5A and 5C). Weaker membrane association of the AH at the ER could expose its serine cluster to increase its turnover rate at the ER. Alternatively, the AH may not engage ER membranes in the context of the full-length protein through an unknown mechanism (Figures 5A and 5B) or DAG-mediated signaling (Eichmann and Lass, 2015) may recruit a factor that promotes association of the AH to the INM and Ser-cluster shielding. Any of these possibilities supports the idea that the AH of Sun2 is a key determinant that promotes stable accumulation of Sun2 at the INM.

**Figure 5:**
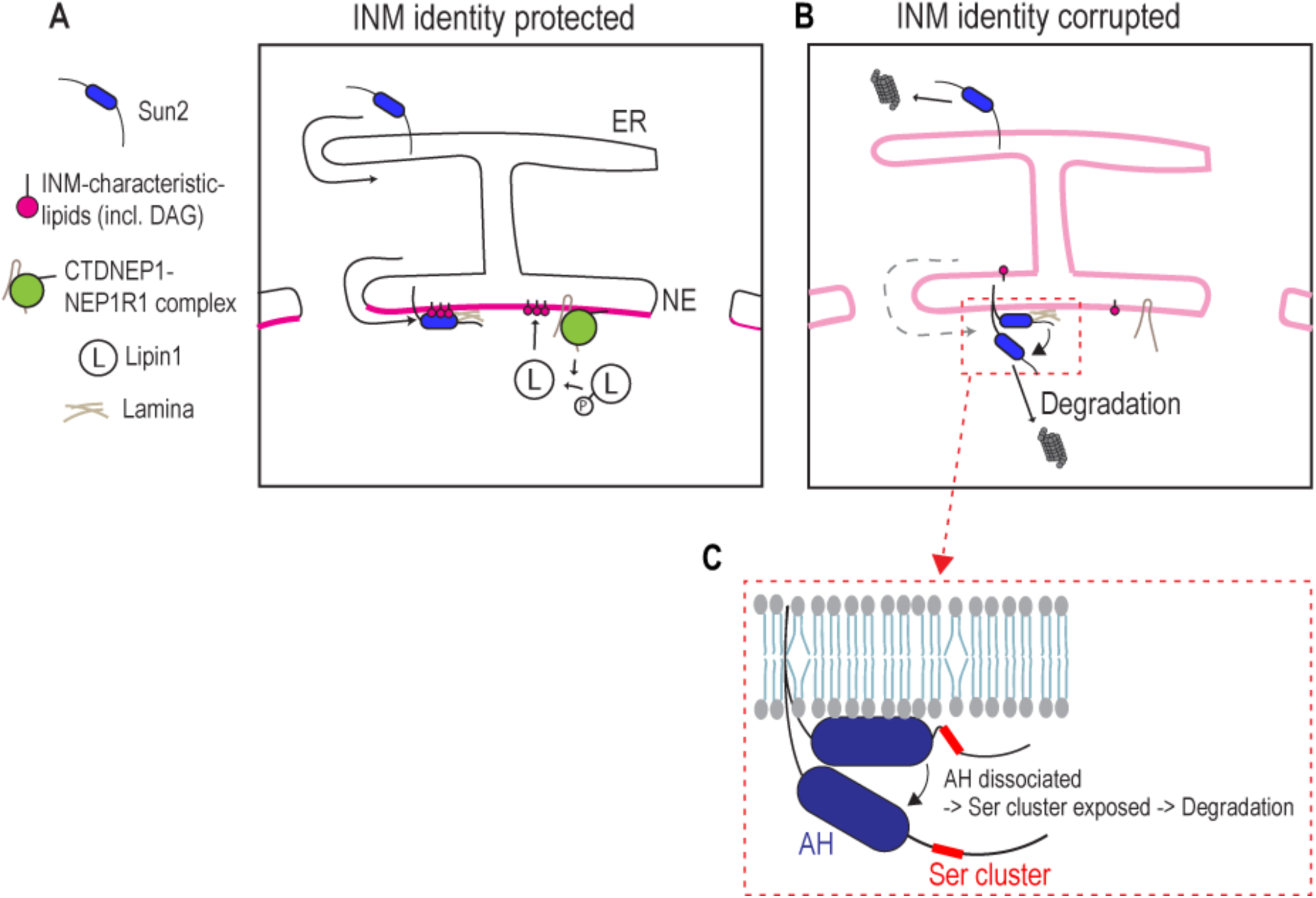
Sensing of a unique INM lipid environment via a membrane binding helix in Sun2 protects it from proteasomal degradation. (A) The lipid identity of the INM depends on CTDNEP1/NEP1R1 to locally regulate DAG levels via lipin 1. The amphipathic helix (blue) in the nucleoplasmic domain of Sun2 promotes its accumulation at the INM by binding directly to INM lipids enriched in DAG to protect it from proteasome degradation. (B and C) Alterations in the INM lipid environment leads to uniform lipid distributions and thus bulk membrane properties. Weak association of the Sun2 AH to membrane lipids exposes upstream serine residues necessary to target Sun2 to proteasomal degradation.

We developed an INM-DAG biosensor that revealed that the INM of animal cells harbors a unique lipid environment. We show that INM DAG levels are maintained by CTDNEP1-mediated control of lipin 1. Disruption of the INM lipid environment, as revealed by reduced enrichment of the INM DAG biosensor, causes rapid degradation of Sun2 demonstrating the importance of the lipid identity of the INM in controlling the INM proteome. Most amphipathic helices are known to sense bulk membrane properties rather than specific lipid species (Drin and Antonny, 2010; Giménez-Andrés et al., 2018; Horchani et al., 2014) and so it is unlikely that the AH of Sun2 binds specifically to DAG. Furthermore, loss of the INM DAG sensor in CTDNEP1 KO cells may not only reflect the lower concentration of DAG at the INM but also overall changes in bulk properties of the INM that can be sensed by the AH of Sun2. Indeed, our data suggest that the AH of Sun2 binds directly to lipids and prefers membrane bilayers with greater packing defects (see Figure S5). Thus, we prefer a model that DAG enrichment at the INM generates a bulk membrane property (e.g. lipid packing defect) (Vamparys et al., 2013; Vanni et al., 2014) that promotes the insertion of the hydrophobic residues of the Sun2 AH. The lack of specificity of the AH of Sun2 towards a specific lipid species leads to its promiscuous binding to the ER and INM, which can be refined by maintaining a more favorable INM environment coupled to additional targeting mechanisms (such as an NLS) (Turgay et al., 2010). The fact that the AH with an NLS promiscuously associates with the ER in CTDNEP1 KO cells suggests that this condition generates uniform bulk membrane properties in both the ER and INM leading to the AH losing its strong association with the INM (Figure 5B). Future work that tests the membrane properties preferred by the AH of Sun2 should help illuminate the distinct nature of the INM lipid environment.

A membrane dissociation and degradation mechanism may also control Sun2 levels in response to external or mechanical signals. Cyclic tensile stress on cells, which is known to lead to reduced Sun2 levels (Gilbert et al., 2019), may affect the bulk properties of the INM to disrupt membrane association of Sun2, leading to its regulated degradation; this in turn may protect NE and genome integrity by reducing cytoskeletal forces imposed on the NE (Gilbert et al., 2019). A Sun2 trimer is required for its binding to KASH proteins (Nie et al., 2016; Sosa et al., 2012; Zhou et al., 2012). Sun2 can also exist as a monomer and in an autoinhibited state, indicating that several mechanisms are in place to ensure Sun2 does not inappropriately engage the cytoskeleton (Nie et al., 2016). The membrane dissociation and degradation mechanism that we describe provides a way to irreversibly prevent Sun2 from inappropriately engaging the cytoskeleton. The detailed mechanistic relationship between the different targeting sequences of Sun2 as well as a physiological context in which Sun2 is degraded in response to changes in lipid metabolism are exciting future directions.

Our data combined with the work of others show that the INM harbors a unique lipid composition across multiple organisms. A common theme is a role for CTDNEP1 in NE-dependent processes including NE sealing, NPC biogenesis, nuclear size regulation, NE breakdown, and nuclear positioning (Bahmanyar et al., 2014; Calero-Cuenca et al., 2021; Jacquemyn et al., 2021; Mall et al., 2012; Mauro et al., 2021; Penfield et al., 2020). These processes may require maintenance of the INM lipid environment by CTDNEP1. Our work links INM lipids to Sun2 levels, thus providing a plausible explanation for why CTDNEP1 is necessary for nuclear positioning via the actin cytoskeleton (Calero-Cuenca et al., 2021) in which Sun2 is also required (Luxton et al., 2010; Zhu et al., 2017). Interestingly, Nem1, the CTDNEP1 homologue in budding yeast, genetically interacts with Mps3, a yeast SUN-domain protein, and mutations in *mps3* result in altered lipid metabolism (Friederichs et al., 2012; Friederichs et al., 2011). Thus, crosstalk between nucleo-cytoskletal linkage and INM lipids may be evolutionarily conserved, although the precise mechanism of Mps3 interaction with membrane lipids may differ from that of Sun2. Determining the direct relationship between CTDNEP1 regulation of lipids at the INM and the phenotypic consequences of disrupting INM lipids will further unveil the role for lipids in INM functions and NE dynamics. Proteomics studies have identified nearly 100 proteins that associate with the NE (Cheng et al., 2019; Korfali et al., 2010; Korfali et al., 2012; Schirmer et al., 2003; Wilkie et al., 2011). Determining the repertoire of INM proteins that respond to local lipid metabolism through direct lipid binding as exemplified by Sun2 is an important future direction. Furthermore, future work that determines mechanisms that control the distribution of specific lipid species at the INM, especially given its continuity with the continuous ER, may reveal how this specialized domain of the ER is established and defined.

## Acknowledgements

We thank M. Hochstrasser, J.M. Gendron, and C. Schlieker (Yale University) for helpful discussions; I. Mérida (CNB), H. Y. Mak (Hong Kong University of Science and Technology), T. Fujimoto (Juntendo University), T. Niki (RIKEN), H. Arai (University of Tokyo), and A. Frost (UCSF), A. Maryniak and A. Holland (Johns Hopkins University), C. Schlieker (Yale University), and Topher Carroll (former Yale School of Medicine) for reagents. We thank E. Rodriguez, M. King and P. Lusk (Yale School of Medicine) for distributing reagents; S. Chou (Yale University) for technical support on protein purification; K. Nelson (Yale University) for the support on flow cytometry; Yale Nucleus Club and BB Club for helpful discussions. This work was supported by: NIH R01 (GM131004) to S.B. Additional support is by Anderson Postdoctoral Fellowship to S.L., NIH (T32s GM100884 and GM007499) and the Gruber Foundation to H.M., NIH (T32 GM722345) to J.W.C.R.

## Author contributions

S.L. and S.B. conceived the project. S.L. performed the majority of the experiments and data analysis. H.M. generated and analyzed the CTDNEP1(EN)-EGFP cell line. J.W.C.R. performed split-cherry experiments and provided data that guided the DAG sensor experiments. S.L. wrote the IJ Macro and Python scripts. S.L. and S.B. wrote the manuscript with input from other authors. S.B. supervised the project.

---

**Figure S1, related to Figure 1.**
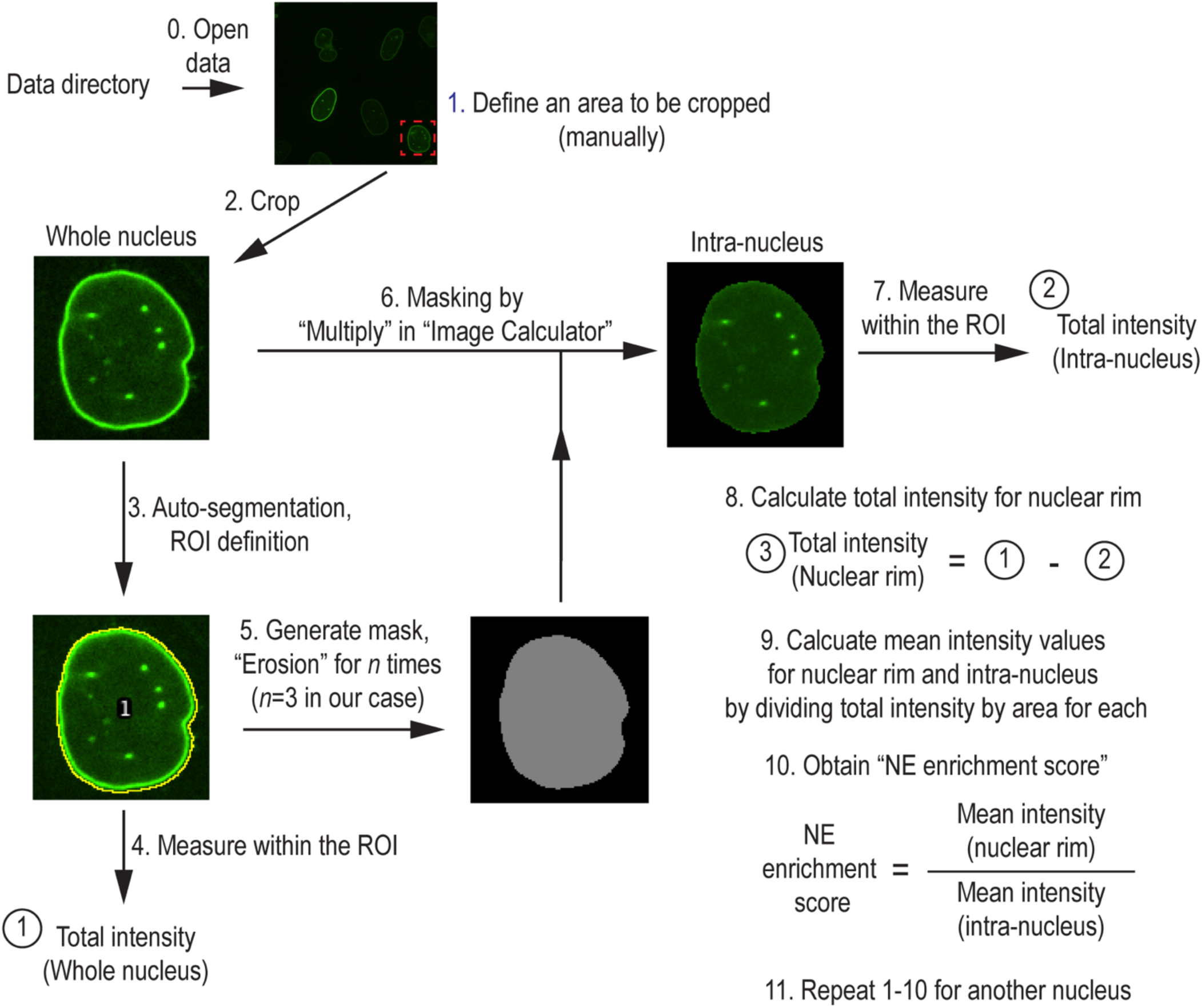
Schematic of the measurement process of the NE enrichment score.

**Figure S2, related to Figure 2A-D.**
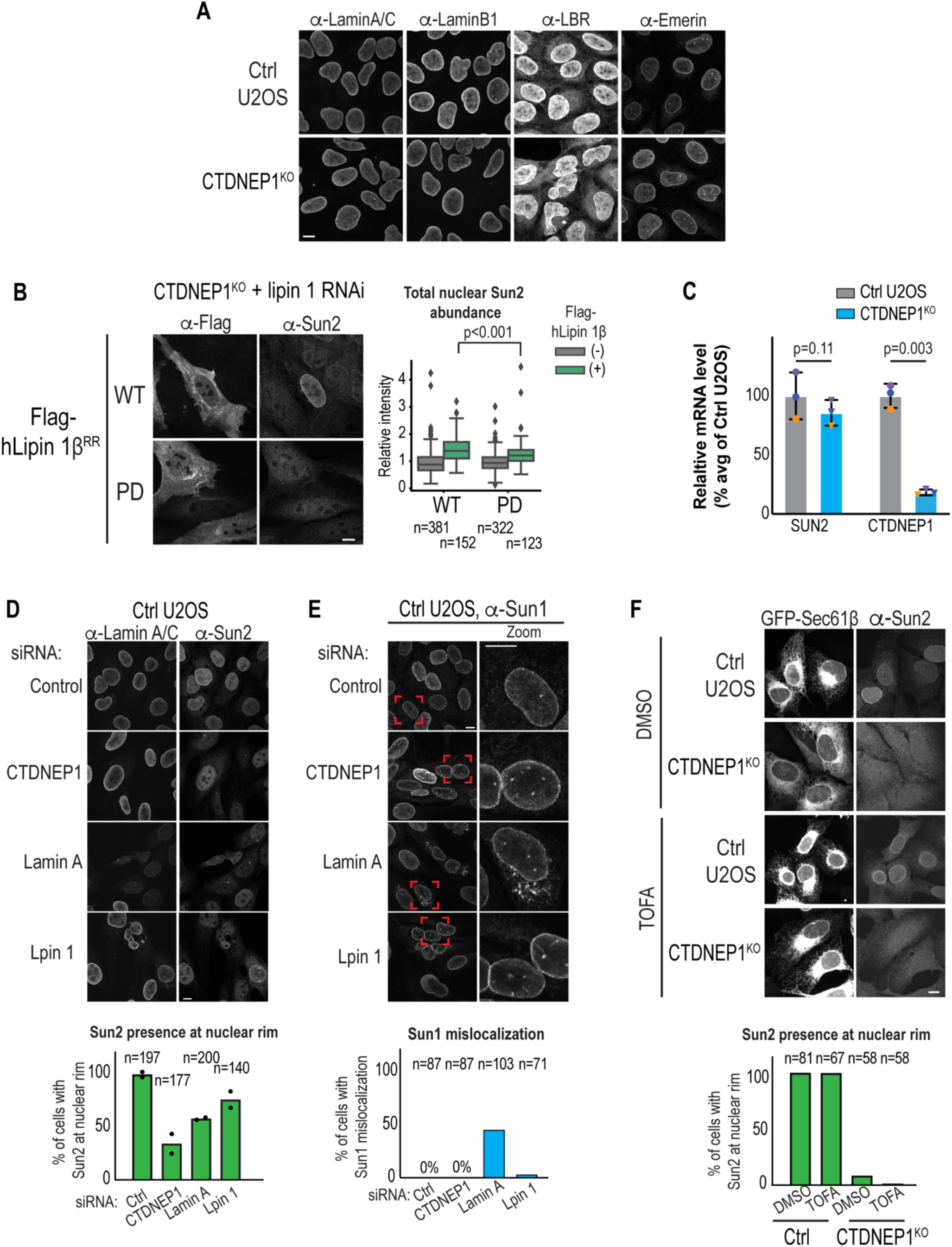
(A) Immunofluorescence of the indicated proteins of U2OS cells. (B) Immunofluorescence of U2OS CTDNEP1 KO cells expressing indicated constructs. Box plot, quantification of Sun2 staining intensity in nuclear area (see methods), pooled from 2 independent experiments. p values, Tukey HSD test. (C) qRT-PCR of indicated cells for indicated genes, shown as fold change in expression relative to mean control values. N = 3 biological replicates. Each pair of replicates is color-coded. p values, paired t test. (D and E) Immunofluorescence U2OS cells treated with indicated siRNAs. (F) Max projections images and quantification of Sun2 localization at nuclear rim of indicated cells transiently expressing GFP-Sec61β and treated with TOFA. Scale bars, 10 µm.

**Figure S3, related to Figure 2E-2G.**
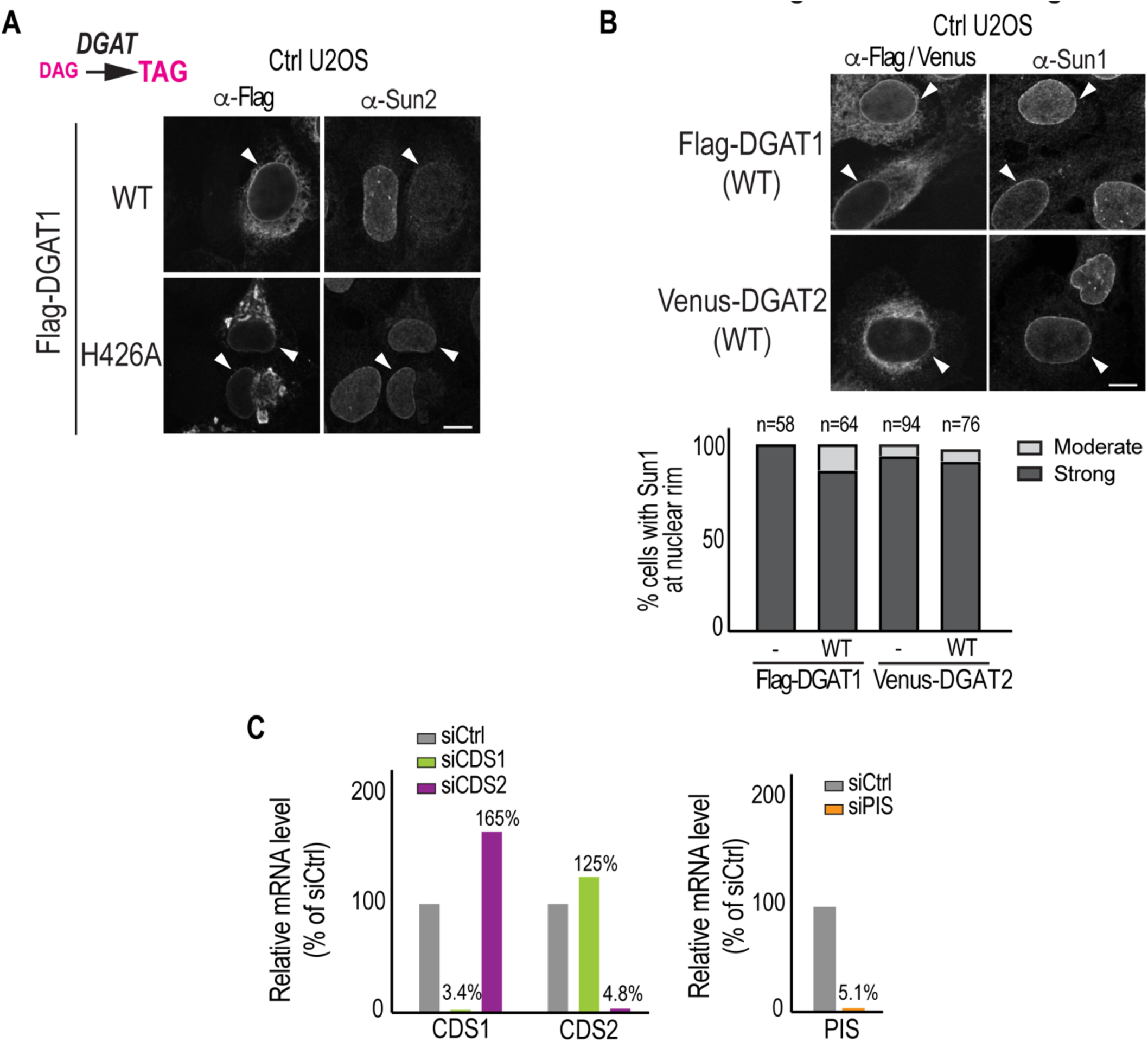
(A) Immunofluorescence of U2OS cells transiently expressing indicated constructs. (B) Representative confocal images of Figure 2F. (C) qRT-PCR of indicated cells for genes indicated, shown as fold change in expression relative to control values. Scale bars, 10 µm.

**Figure S4, related to Figure 3.**
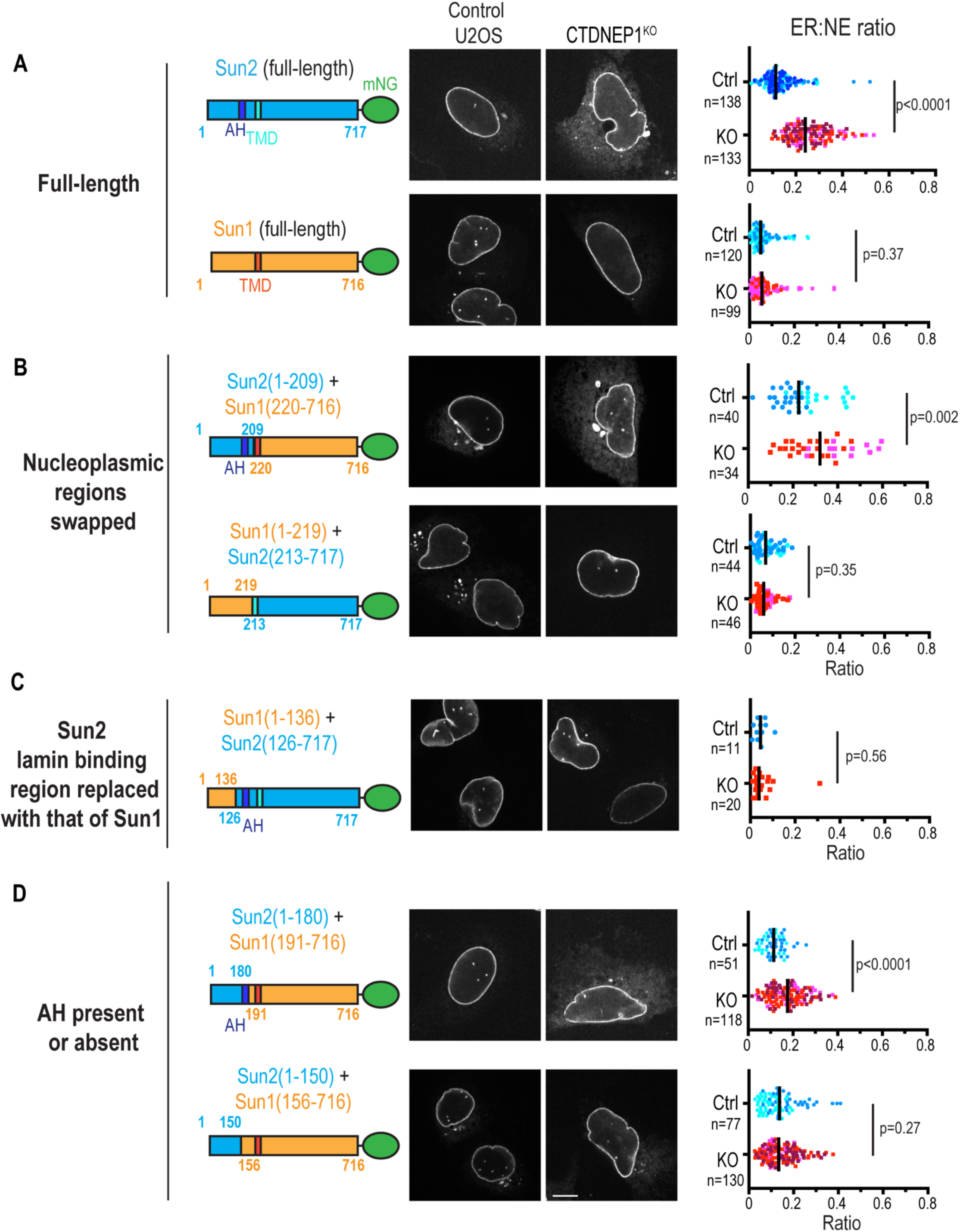
Left: schematic of the Sun2/Sun1 full-length and chimera constructs. Numbers indicate amino-acid residue positions. Middle: representative images of live cells transiently expressing indicated constructs. Scale bar, 10 µm. Right: Plots of ER:NE ratio. Dots are color-coded according to experimental replicates. Bar indicates median. p values, Welch’s t test (for Sun2(1-180)-Sun1(191-716)) or unpaired t test (for others).

**Figure S5, related to Figure 3.**
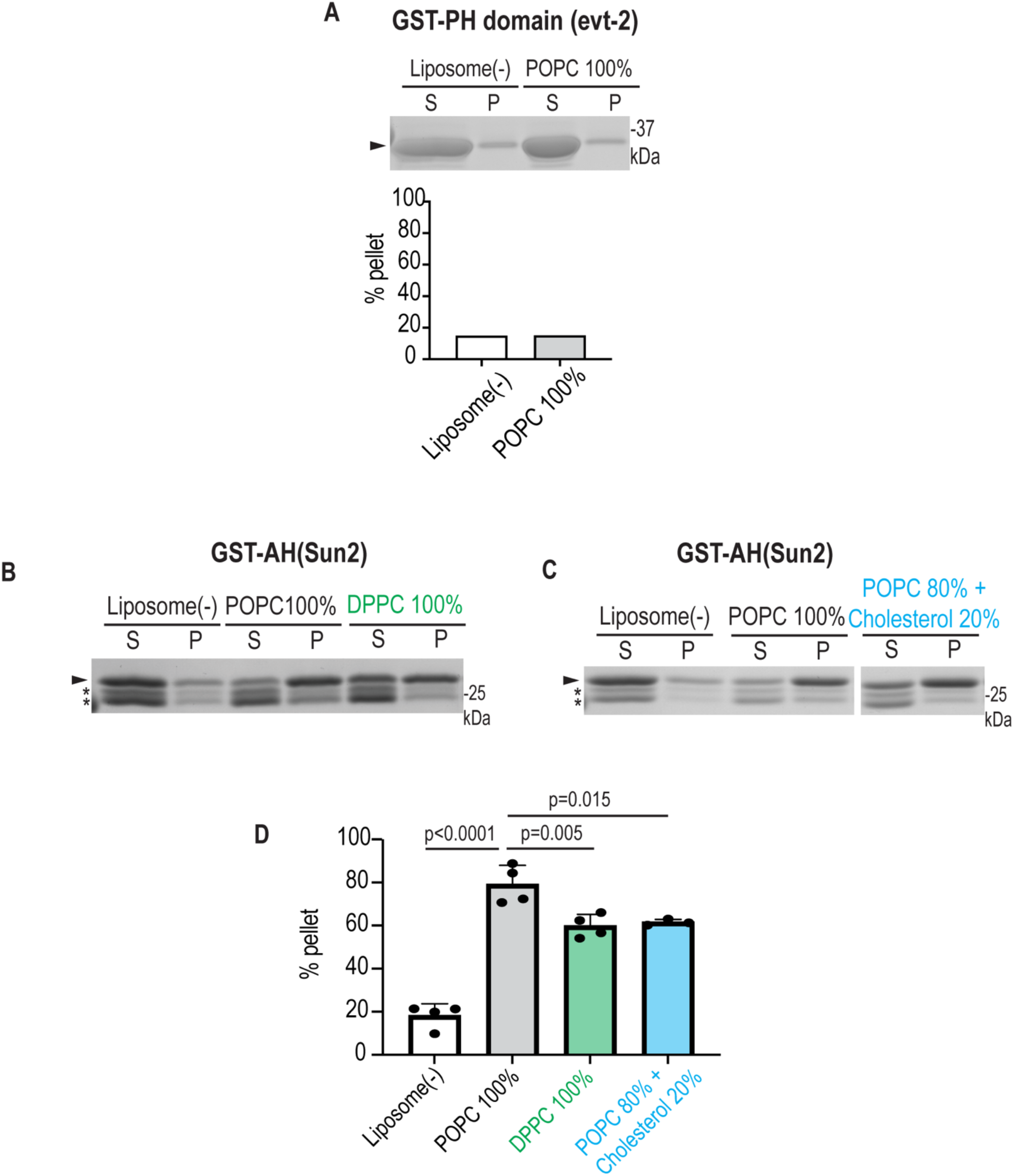
Liposome-cosedimentation with recombinant GST-tagged proteins. (A) GST-tagged PH domain (evt-2) and 1-palmitoyl-2-oleoyl-sn-glycero-3-phosphocholine (POPC) liposome. Plot, percent of protein found in pellet as a ratio of soluble fraction. (B-D) GST-tagged AH peptide (residues 151-180) of Sun2 and liposomes with POPC, dipalmitoyl-sn-glycerol-3-phosphocholine (DPPC) or POPC containing 20 mol% cholesterol. Plot, percent of protein found in pellet as a ratio of soluble fraction. Plots of liposome(-) and POPC 100% are replica of Figure 3B. Three or four experimental replicates. p values, ordinary one-way ANOVA followed by Tukey’s multiple comparisons test. Arrowhead, GST-PH or -AH; Asterisk, purification biproduct.

**Figure S6, related to Figure 3.**
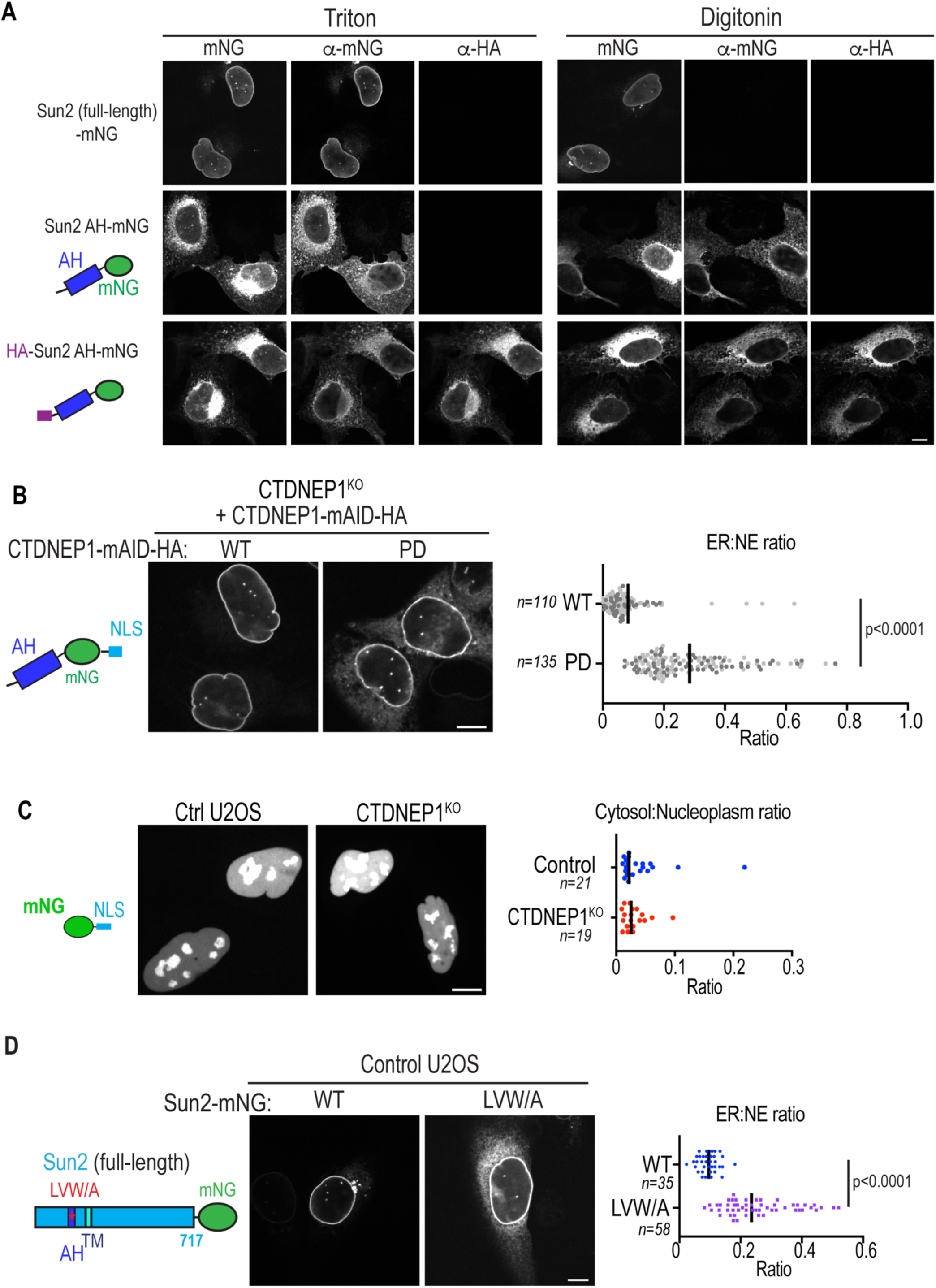
(A) U2OS cells transiently expressing indicated constructs, stained by anti-mNG and anti-HA tag after permeabilized with indicated detergents. (B) Live CTDNEP1 KO cells expressing AH-mNG-NLS and CTDNEP1-mAID-HA WT or D67E (PD) mutant. Plots, ER:NE ratio. Dots are color-coded according to experimental replicates. (C) Live U2OS cells expressing mNG-NLS. Plot, cytosol:nucleoplasm ratio of mNG intensity. (D) Control U2OS cells transiently expressing indicated constructs. Plots, ER:NE ratio. Bars in plots indicate median. p values, Welch’s t test. Scale bars, 10 µm.

**Figure S7, related to Figure 4.**
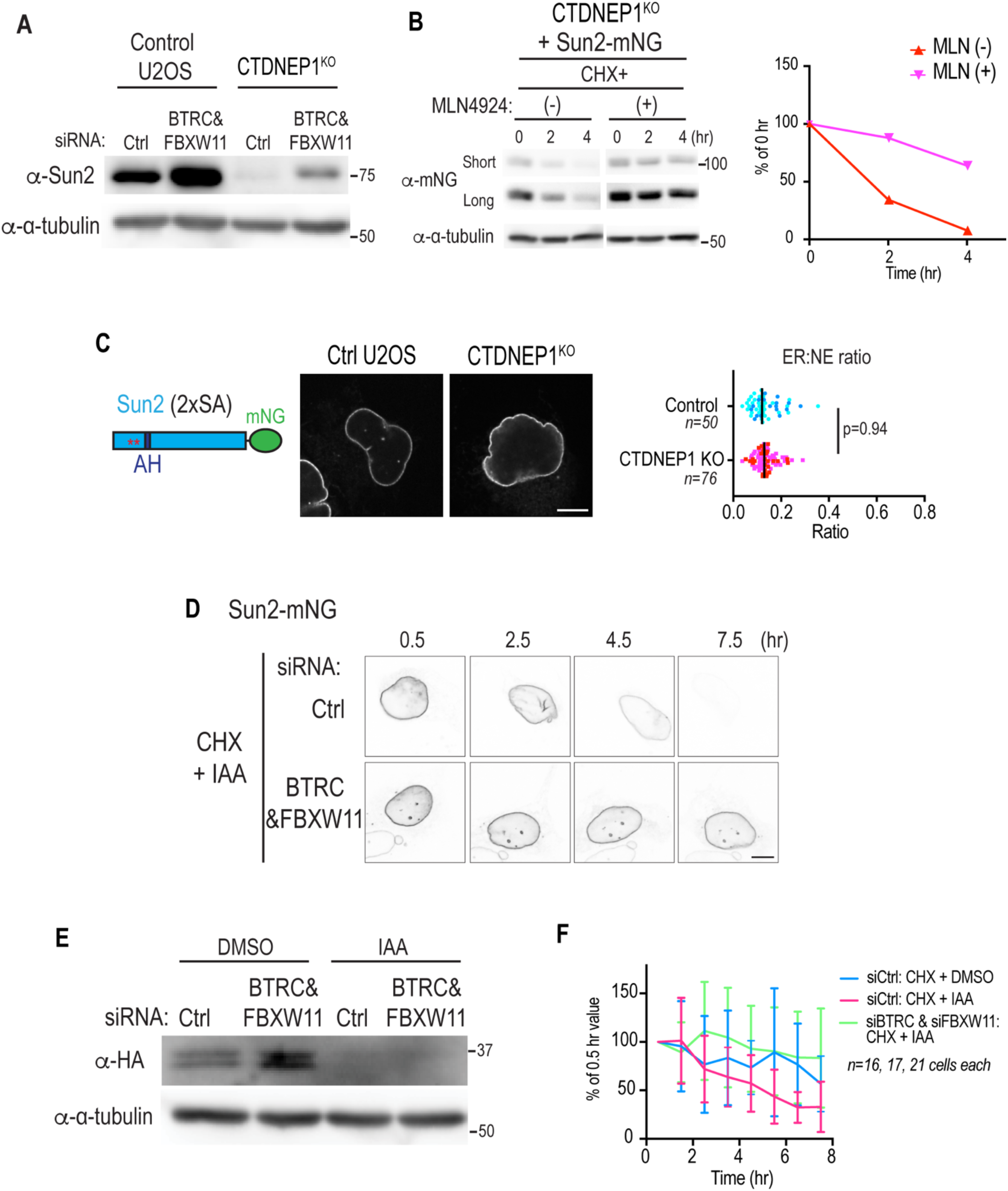
(A) Immunoblot of indicated cells treated with indicated siRNA. (B) Cycloheximide-chase assay of CTDNEP1 KO U2OS cells expressing Sun2 wild-type. MLN4924 was added along with cycloheximide. (C) Live cells expressing Sun2-mNG 2xSA mutant. Plots, ER:NE ratio. Dots are color-coded according to experimental replicates. p value, unpaired t test. (D) Representative time-lapse images of Figure 4I. (E) Immunoblot of DLD-1 OsTIR-1 CTDNEP1(EN)-mAID-HA cells treated with indicated siRNAs and then with IAA for 2 hr. (F) Time-lapse confocal live imaging of DLD-1 OsTIR-1 CTDNEP1EN-mAID-HA cells transiently expressing Sun2-mNG, treated as indicated. mNG intensity in ER region is shown as a percent of the value at 0.5 hr. Data were pooled from 2 independent experiments. Scale bars, 10 µm.

## STAR Methods

### RESOURCE AVAILABILITY

#### Lead Contact

Requests for further information, resources, and reagents should be directed to Shirin Bahmanyar.

#### Materials Availability

Materials generated in this study are available upon request to the Lead Contact.

#### Data and Code Availability

Raw data generated in this study are available upon request to the Lead Contact. ImageJ Macro and Python codes are found at GitHub (https://github.com/shokenlee/Lee_2022_Sun2-Lipid).

#### Key Resource Table

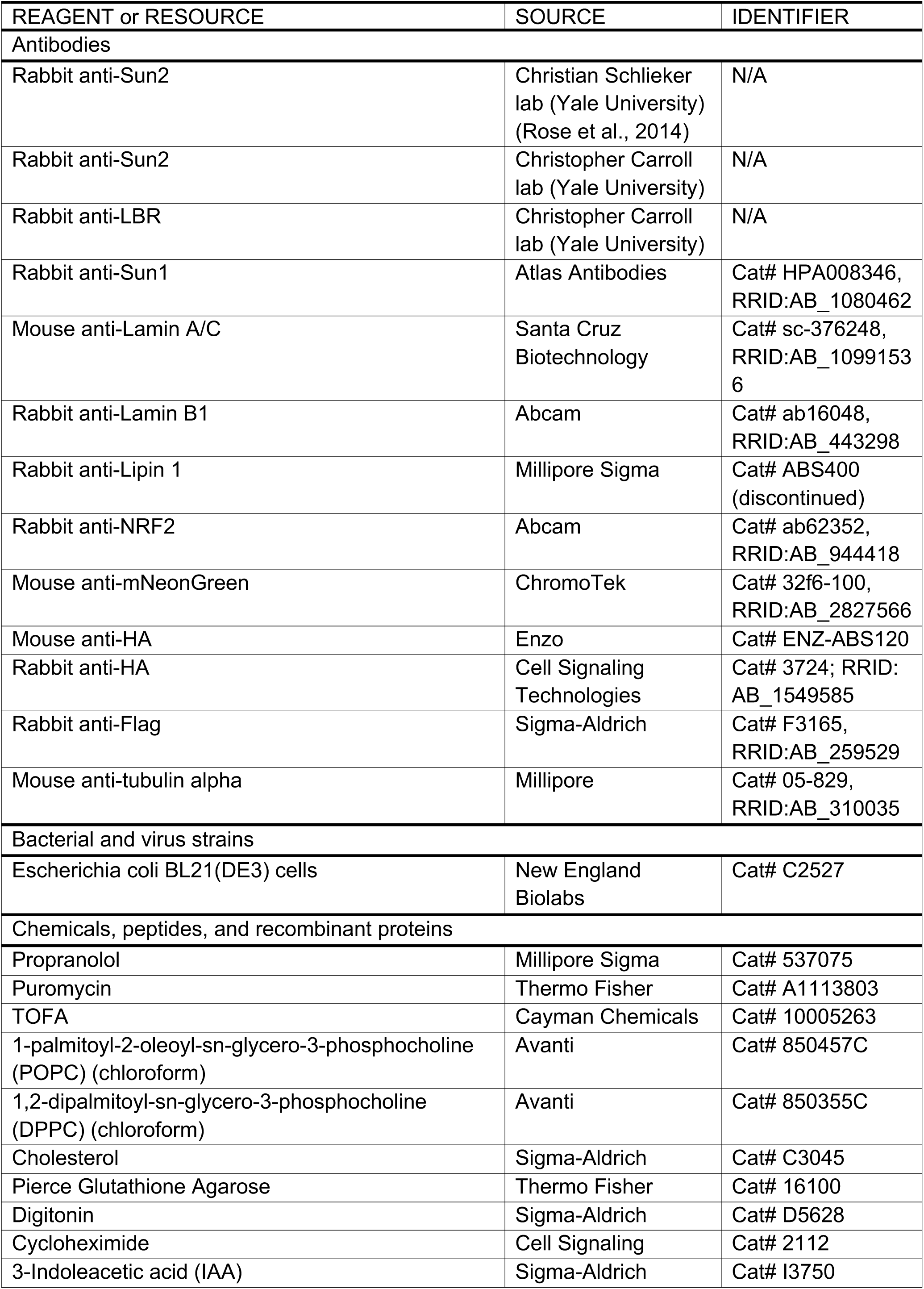
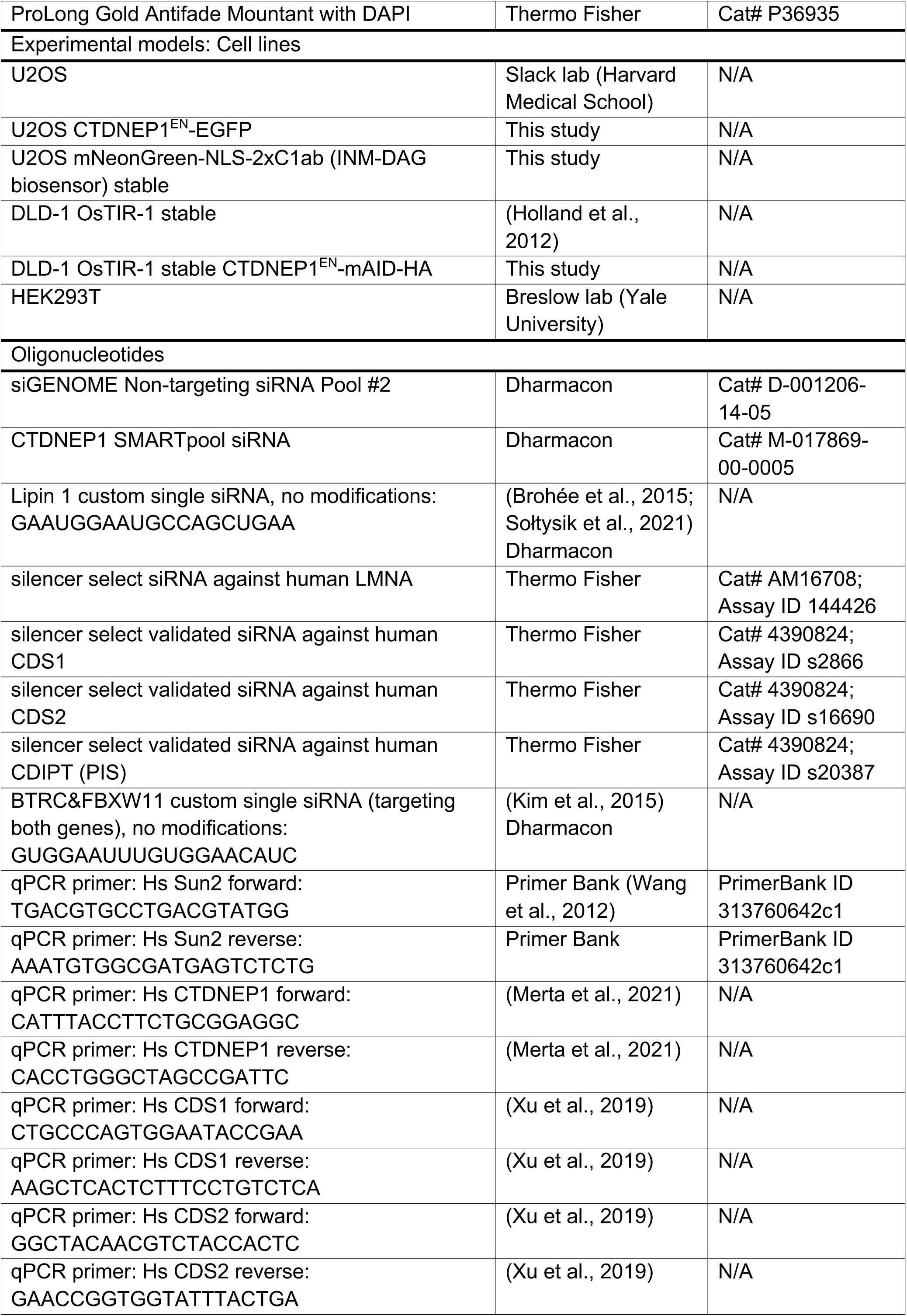
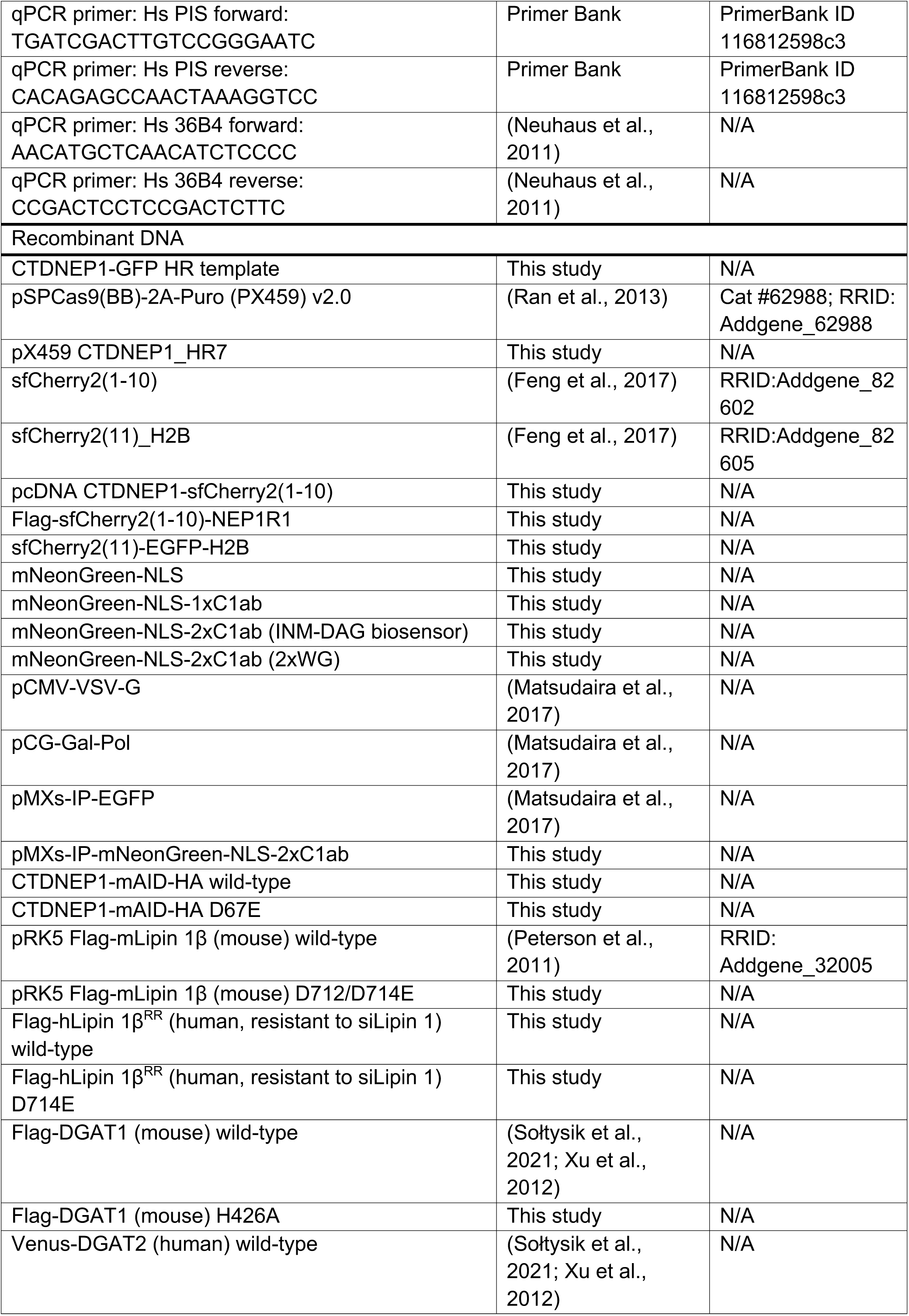
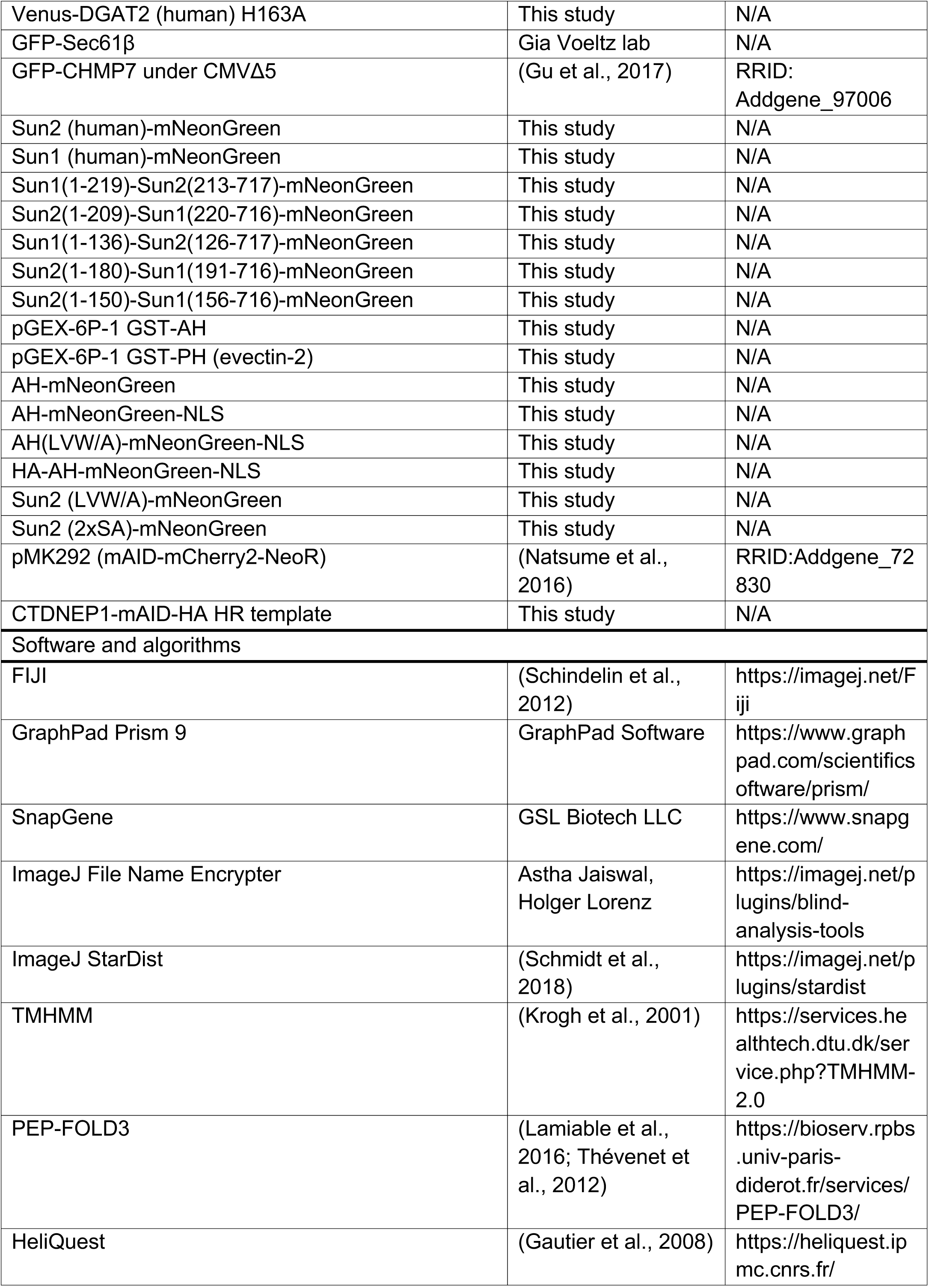
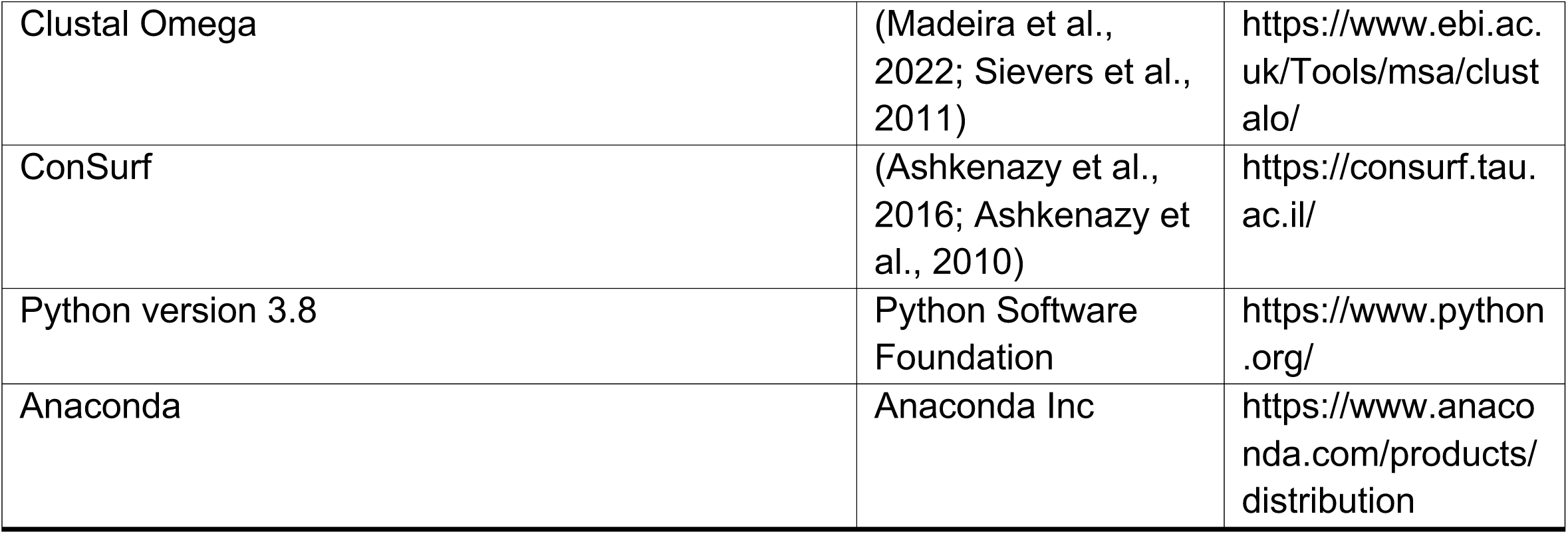

### EXPERIMENTAL MODEL AND SUBJECT DETAILS

#### Plasmid construction

##### General note

Insertion of gene sequence was conducted either by using restriction enzymes from New England Biolabs, Gibson Assembly (New England Biolabs E5510S) or In-Fusion HD Cloning Plus (Takara 638909). Site-directed mutagenesis was performed either by whole-plasmid PCR followed by circularization or by QuikChange Multi Site-Directed Mutagenesis Kit (Agilent 200515). Successful cloning was confirmed by sequencing for all constructs. A bipartite NLS from *Xenopus laevis* nucleoplasmin (KRPAATKKAGQAKKKK) (Dingwall et al., 1988; Robbins et al., 1991; Sołtysik et al., 2019) was used.

##### INM-DAG biosensor

mNG-NLS was first made by inserting mNG ((Shaner et al., 2013); cDNA was a gift from Hiroyuki Arai) followed by NLS in place of EGFP in pEGFP-C2. mNG-NLS-1xC1ab was generated by inserting the coding sequence of C1ab domains (a gift from Isabel Mérida), which corresponds to residues 157-284 of human protein kinase C θ (NP_001310194.1; (Carrasco and Merida, 2004)). mNG-NLS-2xC1ab (INM-DAG biosensor) was generated by inserting another C1ab domain sequence after the first one leaving SRPVLC linker residues in between. Tryptophan mutant was generated by mutating the tryptophans in both C1b domains to glycines, which correspond to W253 in full-length human protein kinase C θ.

##### Sun2-/Sun1-mNG and their chimeras and mutants

The coding sequence for human full-length Sun2 (a gift from M. King and P. Lusk; (Chalfant et al., 2019; May and Carroll, 2018); NP_001186509.1) followed by mNG sequence was subcloned under a clipped CMV promoter (CMVΔ5; (Morita et al., 2012)) by replacing GFP-CHMP7 coding sequence in CMVΔ5-GFP-CHMP7 vector. Sun1 cDNA (716 aa) was isolated by PCR from total cDNA of control U2OS cell line, which was prepared by reverse transcription from total RNA using the iScript Reverse Transcription Supermix (Bio-Rad 1708840). The isolated Sun1 isoform was found to lack the exons 4 and 5 in comparison with the canonical isoform (NM_001130965.3; UniProt ID O94901-8) and was not found in RefSeq proteins of human Sun1 in NCBI Gene database (last search on May 31, 2022), thus appeared to be a novel isoform. The cDNA was inserted between CMVΔ5 and mNG to make Sun1-mNG. Sun2-Sun1 chimeras were generated by fusing the indicated portions from Sun2 and Sun1 under CMVΔ5 promoter by using In-Fusion HD Cloning Plus (Takara). AH-mNG was first generated by replacing the full-length Sun2 in Sun2-mNG vector with AH sequence (Sun2 aa 151-180), and then NLS was inserted after mNG to generate AH-mNG-NLS. Sun2 LVW/A mutant was generated by alanine substitution of L155, V159, L165, L166, W167 and V169 of Sun2. Sun2 2xSA mutant was generated by alanine substitution of S132 and S136.

##### Flag-hLipin 1

Flag-hLipin 1β^RR^ (wild-type or D714E) was subcloned from hLipin 1β^RR^-EGFP constructs (gift from Toyoshi Fujimoto; (Sołtysik et al., 2021)).

#### Mammalian cell lines

U2OS, DLD-1 and HEK293 cells were grown at 37°C in 5% CO2 in DMEM low glucose (Gibco 11885) supplemented with 10% heat inactivated FBS (F4135) and 1% antibiotic-antimycotic (Gibco 15240112). U2OS mNG-NLS-2xC1ab stable cells were grown in the media supplemented with 1 µg/mL puromycin. Cells were used for experiments before reaching passage 25. Cells were tested for mycoplasma upon initial thaw and generation of new cell lines (Southern Biotech 13100-01), and untreated cells were continuously profiled for contamination by assessment of extranuclear DAPI staining.

#### Plasmid and siRNA transfection

For both plasmid and siRNA transfection, cells were seeded to reach 50-80% density on the day of transfection. DNA transfections were performed with Lipofectamine 2000 (Thermo Fisher Scientific 11668) in Opti-MEM (Gibco 31985) with DNA concentrations ranging from 50-300 ng DNA per cm^2^ of growth surface. Cells were imaged or processed after 48 (for Flag-Lipin 1 (mouse and human), Flag-DGAT1, Venus-DGAT2) or 24 hr (others).

RNAi was performed using Dharmafect 1 (Horizon Discovery T-2001) or Lipofectamine RNAiMAX (Thermo Fisher Scientific 13778100) in Opti-MEM according to the manufacturer’s protocol with final siRNA concentration at 20 nM. Cell media was changed on the next day of transfection. When confluent, cells were replated 48 hours after transfection. Cells were processed 72 hours after transfection.

For co-transfection of siRNA and plasmids, siRNA transfection was first performed and media was changed 6 hours after transfection. On the next day, plasmid transfection was performed. Cells were processed 48 hours after plasmid transfection.

#### CRISPR/Cas9 genome editing for generating knockin cell lines

##### U2OS CTDNEP1^EN^-EGFP cell line

Guide RNA sequences were designed using the online CRISPR tool http://crispr.mit.edu and reported no off-target matches: TGGGATGCCGTCTGATGCCC. The guide RNA sequences were synthesized as two oligos with BbsI overhangs and an additional guanidine base 5’ to the protospacer sequence, and the oligos were cloned into pSPCas9(BB)-2A-Puro (PX459) v2.0 into BbsI site. This vector is hereafter called pX459 CTDNEP1_HR. Homology repair template (CTDNEP1-6xGly-EGFP HR) that harbors 6xGly linker sequence before EGFP and 800 bp-homology arms was generated from pEGFP-N1.

U2OS cells were transfected with pX459 CTDNEP1_HR and CTDNEP1-6xGly-EGFP HR vectors with lipofectamine 2000, then treated with 3 µg/mL puromycin for 48 hrs. The remaining cells were grown up and then fluorescent cells were sorted using BD FACSAria. Sorted cells were plated sparsely into a 10 cm culture dish. Cell colonies were trypsinized and picked up with sterile filter paper disks and further grown up in 24 well plates. DNA was extracted from each clone using QiaAmp mini kit, and genotyping PCR and sequencing were performed to select clones that lacked wild-type alleles and had the integration of EGFP into CTDNEP1 loci.

##### DLD-1 OsTIR-1 CTDNEP1^EN^-mAID-HA cell line

The same guide RNA sequence was used as the one used for U2OS CTDNEP1^EN^-EGFP cell line. Vector transfection and colony selection was performed according to (Yesbolatova et al., 2019) with modifications. Briefly, the guide RNA-PX459 vector and CTDNEP1-mAID-HA HR template were transfected into DLD-1 OsTIR-1 stable cells using FuGENE HD (Promega E2311). One day after transfection, cells were collected by trypsinization. Then, collected cells were diluted in media containing final 700 µg/mL G418 and plated in a 10 cm dish. Colonies were trypsinized and picked with filter paper disks, transferred to 96 well plates. Cells were further grown up and duplicated to two 24 well plates and grown until reaching sub-confluency, then either frozen using Bambanker DIRECT medium (Nippon Genetics CS-06-001), or collected with DirectPCR working solution (0.5× DirectPCR Lysis Reagent-Cell [Viagen Biotech, 302-C] containing 0.5 mg/ml of Proteinase K). DirectPCR solution was incubated at 55°C for 6 h then at 85°C for 1.5 h to inactivate Proteinase K. PCR was performed with CloneAmp for genotyping or sequencing.

#### Generation of mNG-NLS-2xC1ab (INM-DAG biosensor) stable cell lines

U2OS mNG-NLS-2xC1ab stable cell lines were generated by retroviral transduction. Retroviruses were generated by transfecting HEK293T cells with pCG-gag-pol, pCMV-VSVG and pMXs-IP-mNeonGreen-NLS-2xC1ab using Lipofectamine 2000. The retroviruses were used to transduce U2OS cells and cells were selected under 1.0 µg/mL puromycin for 2 weeks. Then, cells with fluorescent levels around median range were sorted by BD FACSAria, and plated sparsely to 10 cm culture dish. Cell colonies were trypsinized and picked up with paper filter disks, and further grown up in 24 well plates. For clones that showed no obvious growth anomaly or morphological alteration compared to control U2OS cells, expression level as well as absence of degraded form of the sensor were checked by western blot against mNG, and the sensor localization at nuclear rim was tested by live imaging.

#### Drug treatment

Drug/compound treatment was done as follows: propranolol: 100 µM for 5 min; TOFA: 10 µM for 24 hr; Cycloheximide: 25 µg/mL; MLN4924: 1 µM for 24 hr; IAA: 500 µM. Vehicle control was prepared by diluting the same amount of DMSO as the drug treatment counterpart.

#### Immunofluorescene

Cells on coverslips were fixed in 4% paraformaldehyde (+ 0.1% glutaraldehyde for analyzing an ER marker such as GFP-Sec61β) in PBS for 15 min at room temperature, permeabilized in 0.1% Triton X-100 for 3 min at room temperature, then blocked in 3% BSA in PBS for 30 min at room temperature. For digitonin permeabilization, cells were permeabilized with 10 µg/mL digitonin on ice for 10 min. Samples were then incubated with primary antibodies in 3% BSA in PBS for 1.5 hours at room temperature or for overnight at 4°C. Samples were washed with PBS 3 times, then incubated with secondary antibodies in 3% BSA in PBS for 45 min at room temperature in the dark with rocking. Samples were then washed with PBS 3 times. Coverslips were mounted with ProLong Gold Antifade reagent with DAPI. Primary antibody concentration: anti-Sun2 (Carroll lab, used only in Figure 2A) 1:1000; anti-Sun2 (Schlieker lab, used in other experiments) 1:1000; anti-LBR 1:250; anti-SUN1 1:250; anti-Lamin A/C 1:1000; anti-Lamin B1 1:1000; anti-mNeonGreen 1:1000; mouse anti-HA 1:1000; anti-Flag 1:1000.

#### Live cell imaging

For live imaging, cells were plated in µ-Slide 8 Well Glass Bottom (ibidi 80827). Samples were imaged in a CO_2_-, temperature-, and humidity-controlled Tokai Hit Stage Top Incubator. The imaging media used was Fluorobrite DMEM (Gibco A1896701) supplemented with 10% FBS, 2 mM L-glutamine (Gibco A2916801) and 1% antibiotic-antimycotic (Gibco 15240112).

For live imaging of Sun2-mNG with cycloheximide, cell media was exchanged for the media containing 25 µg/mL cycloheximide and 500 µM IAA as indicated, then imaging was initiated after 30 min of the media change.

#### Microscopy

Fixed cell imaging and live cell imaging without time lapse were performed on an inverted Nikon Ti microscope equipped with a Yokogawa CSU-X1 confocal scanner unit with solid state 100-mW 488-nm and 50-mW 561-nm lasers, using a 60×1.4 NA plan Apo oil immersion objective lens, and a Hamamatsu ORCA R-2 Digital CCD Camera. Live cell time-lapse imaging was performed on an inverted Nikon Ti Eclipse microscope equipped with a Yokogawa CSU-W1 confocal scanner unit with solid state 100 mW 405, 488, 514, 594, 561, 594 and 640 nm lasers, using a 60x 1.4 NA plan Apo oil immersion objective lens and/or 20x plan Fluor 0.75 NA multi-immersion objective lens, and a prime BSI sCMOS camera.

#### Image analysis

##### NE enrichment score of INM-DAG sensor

Raw image data were semi-automatically measured with a custom ImageJ Macro code (see Figure S1 for a schematic) and measurement results were analyzed with a custom Python code. Firstly, the user was prompted by the macro to count the number of cell nuclei that were amenable to quantification. Cell nuclei with wrinkles or any abnormal shape were not included in quantification because these structural changes tended to blur the boundary between nuclear membrane and nucleoplasm and thus hampered correct measurement. Then, a nucleus on an optimal z frame was manually selected by a rectangle. Following the selection, the macro continued on the following process. The nucleus image was cropped (called “Whole nucleus”), auto-segmented by Li method and the ROI for the whole nucleus was defined. In some cases, in particular where the overall intensity was low, the auto-segmentation failed to find the nucleus and no ROI was defined. Results from those cases were excluded later on the Python code based on area value = 0. To measure intensity from “intra-nucleus” (i.e., nucleus area excluding nuclear rim), firstly a mask image was generated by performing “Erosion” three times of the binary image that was generated from the ROI. This mask was multiplied with “Whole nucleus” image by “Image Calculator” to leave signals only from the intra-nucleus area (called “Intra-nucleus”). Total intensity within the ROI was measured for “Whole-nucleus” and “Intra-nucleus”. Total intensity of the binary mask within the ROI was also measured, which gives a value that is equal to the value of the area of “Intra-nucleus”. The total intensity and area values of the “Nuclear rim” were obtained by subtraction of the values of “Intra nucleus” from “Original nucleus”. Mean value of “Whole nucleus” and “Nuclear rim” were calculated by dividing the total intensity by the area of each. Finally, after subtraction of background mean value, the ratio of “Nuclear rim” to “Intra nucleus” was calculated to give the “NE enrichment score”. The measurement was repeated for all cells in images in a given directory. The measurement results were exported to CSV files and analyzed by Python Pandas, Matplotlib and Seaborn libraries. Statistical analysis was performed by Tukey HSD test using the Python Statsmodels library.

##### Sun2/Sun1 NE localization

Endogenous Sun2/Sun1 NE localization was categorized into three bins: “strong”, “moderate” and “non” based on visual impression of the clarity and brightness of Sun2/Sun1 localization at the nuclear rim. Categorization was performed blindly by using File Name Encrypter in Blind Analysis Tools of Fiji.

##### Sun2 expression intensity in nucleus area

Raw image data were automatically measured with a custom ImageJ Macro code and measurement results were analyzed with a custom Python code. Firstly, images were max-projected and split to DAPI, Flag-tag (absent in case for siCDS/PIS experiments) and Sun2 channels. Then, from DAPI channel, ROIs for nuclear area were defined by StarDist plugin with the default settings. Within the ROIs, mean intensity values of DAPI, tag, and Sun2 as well as area were measured and exported to a CSV file. The resulting CSV files were concatenated and analyzed using Python Pandas, NumPy, Matplotlib and Seaborn libraries. Background was subtracted from Sun2 intensity. In most cases background=2000 was applied. Flag tag-positive cells were determined based on a threshold value, which was determined by examination of several raw images. Typically mean Flag intensity=3000 was applied. Mitotic cells and small DAPI signals including cell debris and micronuclei were excluded from analysis based on relatively high intensity of DAPI and small area size, respectively. After subtraction of background value, Sun2 intensity was normalized against the mean value of cells without tag expression in each transfection condition. Statistical analysis was performed by Tukey HSD test using the Python Statsmodels library.

##### NE enrichment of Sun2/Sun1 and its chimera/mutants (ER:NE ratio)

In a given image on Fiji, a line with 5-pixel width and 5-µm length was drawn from cytoplasm to nucleus, centered on the nuclear rim. The maximum value along the line was considered the “NE” value. The average value of the first 5 pixel along the line (namely 5×5 pixels) was considered the “ER/cytoplasm” value. Average value from the cell-free area was used to subtract background. Then the ER:NE ratio was calculated.

##### Nucleus enrichment of mNG-NLS (cytosol:nucleoplasm ratio)

Mean intensity values of manually-drawn rectangular regions (5×5 pixels) were measured in cytosolic and nucleoplasmic area. When necessary, brightness/contrast was modified to distinguish cytosolic area from non-cell area. Nucleoli was avoided when drawing a nucleoplasm region for measurement. Average value from the cell-free area was used to subtract background. Then the cytosol:nucleoplasm ratio was calculated.

##### Sun2-mNG intensity at NE and ER under IAA treatment

Similarly to NE enrichment analysis, line scan was performed with a line crossing the nuclear rim. The maximum intensity was considered Sun2-mNG intensity at the NE, and the average value of the first 5 pixel along the line (namely 5×5 pixels) was considered the ER value.

#### Western blot

Cells were lysed with ice-cold RIPA buffer (25 mM Tris pH 7.4, 1% NP-40, 0.5% sodium deoxycholate, 0.1% SDS, 150 mM NaCl, and 1 tablet/50 ml cOmplete Mini protease inhibitor cocktail (Roche 11836153001)) with cell scraper, incubated on ice for 15 min, and then centrifuged at >20,000xg (15,000 rpm) for 15 min at 4°C. Protein concentration was determined using the Pierce BCA Protein assay kit (Thermo Scientific 23225). 10-15 μg of whole cell lysates/lane were run on 8-15% polyacrylamide gels dependent on target size, and protein was wet transferred to 0.22 μm nitrocellulose membranes (Bio-Rad 1620112). Membranes were blocked in 5% nonfat dry milk in TBS (for lipin 1) or 1 or 3% BSA in PBS (for targets other than lipin 1) for 30 min. Membranes were then incubated with primary antibodies in milk or BSA for 1.5 hours at room temperature or overnight at 4°C with rocking.

Membranes were washed 3 times for 5 min in TBST, then incubated with goat anti-mouse or rabbit IgG secondary antibodies (Thermo Fisher 31430 or 31460) in 5% milk in TBST for 45 min at room temperature with rocking. Membranes were washed 3 times for 5 min in TBST. Clarity or Clarity Max ECL reagent (Bio-Rad 1705060S, 1705062S) was used to visualize chemiluminescence, and images were taken with a Bio-Rad ChemiDoc Imaging System. Blot was quantified by using Fiji. Antibody concentration was the following: anti-Sun1, Lipin 1, HA, NRF2, 1:1000; anti-Lamin A/C, Sun2 (Carroll lab), mNeonGreen 1:3000; α-tubulin 1:5000; secondary antibodies 1:10000

#### Cycloheximide chase assay

Cell media was exchanged with 25 µg/mL cycloheximide. After the indicated time, cells were collected for western blot. Immunoblot intensity of mNG or HA-tag was quantified and normalized to that of α-tubulin, and is shown as % of the value of 0 hr.

#### Quantitative real-time PCR

RNA was harvested using the RNeasy Mini kit (Qiagen 74104) using the manufacturer’s protocol, using Qiashredders (Qiagen 79654) for tissue homogenization and with additional RNase-free DNase (Qiagen 79254) treatment after the first RW1 wash and subsequently adding another RW1 wash. RNA was eluted with RNAse-free water and diluted to 50 ng/μl. RNA was subject to reverse transcription using the iScript Reverse Transcription Supermix (Bio-Rad 1708840) with 400 ng RNA per reaction. The subsequent cDNA was diluted 1:5 for RT-qPCR. cDNA was analyzed for RT-PCR using the iTaq universal SYBR Green Supermix (Bio-Rad 1725120) with the CFX384 Touch Real-Time PCR Detection System (Bio-Rad). Production of a single amplicon was confirmed by melt curve analysis. Cycle threshold values were analyzed using the ΔΔCt method.

#### Recombinant protein purification

GST-tagged proteins were expressed in *Escherichia coli* BL21(DE3) cells. Cells were grown at 37 °C to an OD600nm of 0.6-0.9 and then cooled at 20 °C. Protein expression was induced with isopropyl βD-1-thiogalactopyranoside (IPTG) at 20 °C for 16 h, and cells were harvested by centrifugation and stored at −80 °C. Frozen cells were resuspended in buffer A (1x PBS, 1 mM EDTA, 1 mM EGTA, pH 8.0) and lysed by sonication. After adding DTT and CHAPS detergent at final 1 mM and 2% w/v concentration, respectively, sample was centrifuged at 20,000 rpm with Beckman Coulter type 70 Ti rotor at 4 °C for 30 min. The supernatant was incubated with pre-equilibrated Pierce Glutathione Agarose at 4 °C for 1 h. The resin was centrifuged at 2,000 rpm at 4 °C for 2 min, washed 3 times with buffer B (50 mM Tris-HCl pH 8.0, 150 mM NaCl, 1 mM DTT, 1% CHAPS) and transferred to pre-wet Econo-Pac Chromatography Columns (Bio-Rad 7321010). Protein was eluted with buffer C (50 mM Tris-HCl pH 8.0, 150 mM NaCl, 10 mM reduced glutathione, 0.7% CHAPS), aliquoted and analyzed by SDS-PAGE and Coomassie blue stain. Fractions containing the protein were diluted 3-fold in buffer C so that the final CHAPS concentration is below its critical micelle concentration, which is 0.4-0.6%. The diluted fractions were dialyzed in 1xPBS using Slide-A-Lyser Dialysis Cassette G2 10,000 MWCO 15 mL (Thermo Fisher 87731). Dialyzed protein solution was concentrated roughly 3-fold by centrifugation with Amicon Ultra-15 Centrifugal Filter Unit (Millipore UFC901024) at 4,000 rpm at 4°C for 15 min, then flash-frozen and stored at −80 °C until used.

#### Liposome co-sedimentation

Lipid mixtures were dried under nitrogen gas and then under vacuum for 1 hr at room temperature. Dried lipids were hydrated liposome-binding buffer (20 mM HEPES-NaOH pH 7.4, 150 mM NaCl, 1 mM MgCl_2_, 1 mM DTT) for 60 min at 37°C, vortexed for 1 min, and subjected to 3 rounds of freeze-thaw cycles with liquid nitrogen. To remove protein aggregates, the protein solution was subjected to centrifugation using a TLA120 fixed angle rotor (Beckman) at 55,000 rpm for 15 min at 4°C before use. 10 µg of proteins were incubated with 50 nmol liposomes for 10 min at room temperature, and the mixture was centrifuged at 55,000 rpm for 30 min at 20°C using a TLA120 fixed angle rotor (Beckman). The resultant supernatant and pellet were subjected to SDS–PAGE, and the proteins and lipids were stained with Coomassie blue. The intensities of individual bands were quantified with Fiji.

#### Secondary structure analysis

Secondary structure analysis was done with PEP-FOLD3. HeliQuest was used to generate a helical wheel projection and to obtain hydrophobic moment <µH> and net charge Z values, which yielded a discriminant factor D = 0.944 (<µH>) + 0.33 (z) as described in HeliQuest (https://heliquest.ipmc.cnrs.fr/HelpProcedure.htm).

#### Conservation score analysis

Amino-acid sequences of putative Sun2 orthologues in jawed vertebrates were obtained from the list of “Orthologs” of human Sun2 in NCBI Gene database. Species were chosen such that they include birds, turtles, alligators, lizards, mammals, amphibians and cartilaginous fishes. The sequence of the zebrafish homologue, which was not found in the list, was obtained from HomoloGene in NCBI database (ID: 9313). Accession numbers of proteins are the following: *Homo sapiens*: NP_001186509.1; *Pan troglodytes* (chimpanzee): XP_003317285.1; *Canis lupus familiaris* (dog): XP_538371.3; *Bos taurus* (cow): NP_001095789.1; *Mus musculus*: NP_001192274.1; *Rattus norvegicus*: XP_235483.6; *Phyllostomus hastatus* (bat): XP_045690878.1; *Gallus gallus* (chicken): XP_416257.4; *Pogona vitticeps* (bearded dragon): XP_020643298.1; *Chelonia mydas* (sea turtle): XP_037737727.1; *Xenopus (Silurana) tropicalis* (frog): XP_002933805.2; *Rhinatrema bivittatum* (caecilian): XP_029446057.1; *Danio rerio* (zebrafish): XP_001919691.1; *Amblyraja radiata* (skate): XP_032869439.1. A multiple sequence alignment and phylogenetic tree were generated using Clustal Omega with the default settings. The conservation score was obtained using the ConSurf server with the default settings by providing the multiple sequence alignment (query: human Sun2) and phylogenetic tree as inputs. Low confidence of the score was presented as yellow color of a residue.

#### Statistical analysis

GraphPad Prism 8 was used for all statistical analysis otherwise specified in methods. Color-coding of each experiment repetition was based on Superplot (Lord et al., 2020). Sample size required for reliable statistical analysis was determined before performing experiments using Sample Size Calculator (https://clincalc.com/stats/samplesize.aspx). Statistical tests used, sample sizes, definitions of n and N, and p values are reported in figures and/or figure legends.

